# Solvent Precipitation SP3 (SP4) enhances recovery for proteomics sample preparation without magnetic beads

**DOI:** 10.1101/2021.09.24.461247

**Authors:** Harvey E. Johnston, Kranthikumar Yadav, Joanna M. Kirkpatrick, George S. Biggs, David Oxley, Holger B. Kramer, Rahul S. Samant

**Affiliations:** Signalling Programme, The Babraham Institute, Cambridge, CB22 3AT, United Kingdom; Mass Spectrometry Facility, The Babraham Institute, Cambridge, CB22 3AT, United Kingdom; Proteomics STP, The Francis Crick Institute, London, NW1 1AT, United Kingdom; GlaxoSmithKline, Gunnels Wood Road, Stevenage, Hertfordshire, SG1 2NY, United Kingdom; Medical Research Council London Institute of Medical Sciences, Imperial College London, Hammersmith Hospital, London, W12 0NN, United Kingdom

**Keywords:** Proteomics, sample preparation, protein precipitation, contaminant removal, LC-MS/MS, microparticles, protein aggregation

## Abstract

Complete, reproducible extraction of protein material is essential for comprehensive and unbiased proteome analyses. A current gold standard is single-pot, solid-phase-enhanced sample preparation (SP3), in which organic solvent and magnetic beads are used to denature and capture proteins, with subsequently washes allowing contaminant removal. However, SP3 is dependent on effective protein immobilisation onto beads, risks losses during wash steps, and experiences a drop-off in protein recovery at higher protein inputs. Magnetic beads may also contaminate samples and instruments, and become costly for larger scale protein preparations. Here, we propose solvent precipitation SP3 (SP4) as an alternative to SP3, omitting magnetic beads and employing brief centrifugation—either with or without low-cost inert glass beads—as the means of aggregated protein capture. SP4 recovered equivalent or greater protein yields for 1–5000 µg preparations and improved reproducibility (median protein *R*^2^ 0.99 (SP4) *vs*. 0.97 (SP3)). Deep proteome profiling (n = 9,076) also demonstrated improved recovery by SP4 and a significant enrichment of membrane and low-solubility proteins vs. SP3. The effectiveness of SP4 was verified in three other labs, each confirming equivalent or improved proteome characterisation over SP3. This work suggests that protein precipitation is the primary mechanism of SP3, and reliance on magnetic beads presents protein losses, especially at higher concentrations and amongst hydrophobic proteins. SP4 represents an efficient and effective alternative to SP3, provides the option to omit beads entirely, and offers virtually unlimited scalability of input and volume—all whilst retaining the speed and universality of SP3.

**BRIEF:** Solvent precipitation SP3 (SP4) captures aggregated protein for proteomics sample clean-up by omitting magnetic beads, instead employing brief centrifugation—with or without low-cost inert glass beads. SP4 offers improvements to protein yields, higher reproducibility, and greater recovery of membrane proteins, with verifications from three labs. Protein precipitation appears to be the primary mechanism of SP3, with reliance on magnetic beads presenting protein losses, especially at higher concentrations. SP4 presents an effective alternative to SP3 with improved scalability and equal speed and universality.

**GRAPHICAL ABSTRACT:** 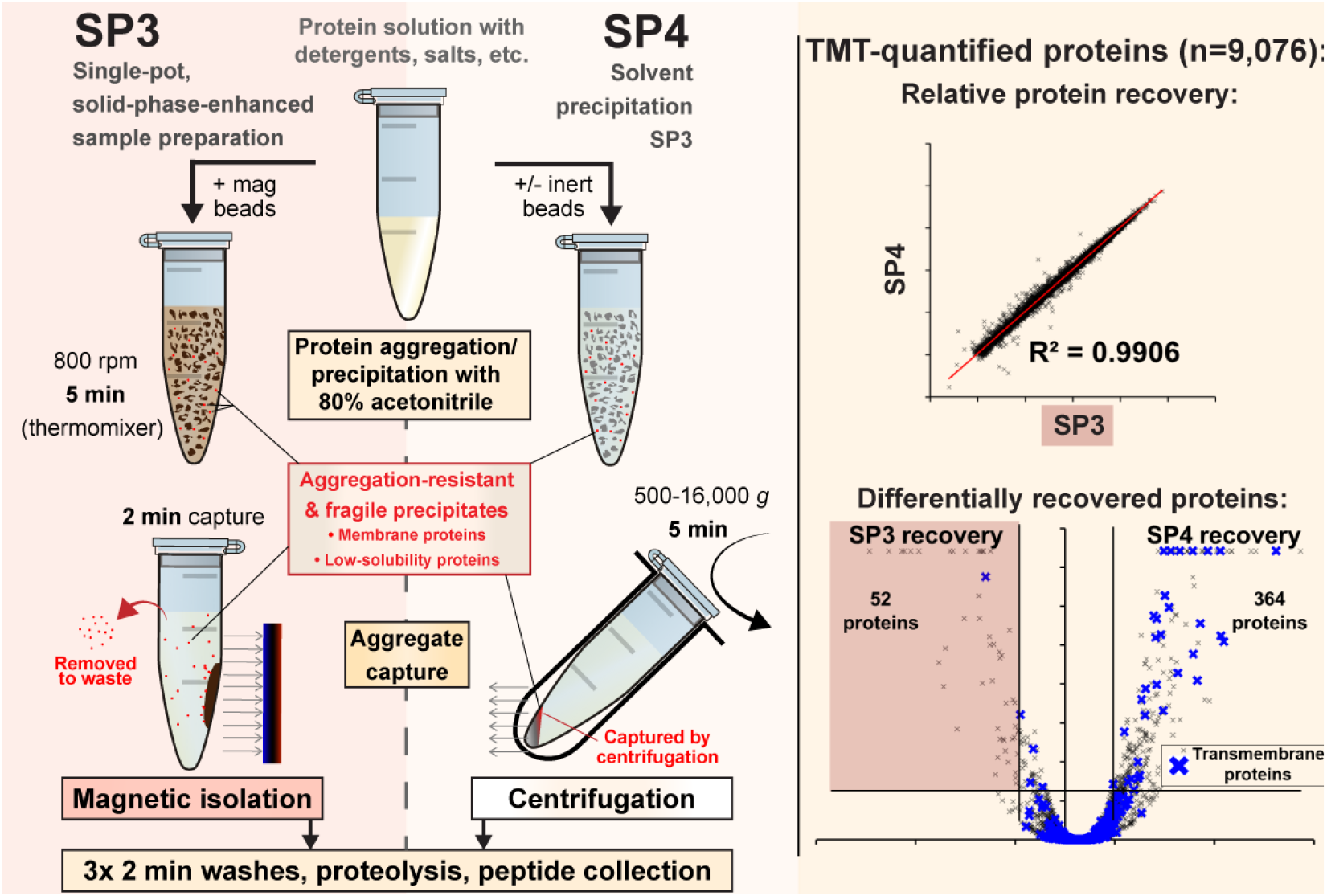

## Introduction

At a fundamental level, a common goal of proteomics experiments is to provide a quantitative snap-shot of all proteins contained within a given cellular sample, at a given time, in as global and unbiased a way as possible (1). Central to achieving this goal is the ability to preserve and isolate proteins from the starting material. Most effective cell lysis buffers often contain high concentrations of detergents or salts: contaminants that substantially interfere with proteome characterisation by mass spectrometry (MS), and must be removed (2). Many methods have been proposed to address protein or peptide clean-up for contaminant removal, which still represents one of the largest sources of sample loss and variability in a proteomics experiment (3).

Generally, clean-up steps are incorporated at two stages of a bottom-up proteomics workflow: protein isolation from the initial cell lysate, and/or peptide isolation after enzyme digestion. The choice of workflow places limitations on the cleanup methods available for use. For example, although detergents provide an effective means of solubilising and isolating proteins from other macromolecules (e.g., lipids and carbohydrates), they interfere with digestion, chromatography and MS, and are incompatible with solid-phase extraction peptide clean-up methods. Various methods for detergent depletion have been developed, each with specific strengths and limitations (4–11).

Protein precipitation provides an effective means of purifying proteins from many forms of contamination (12). By eliminating water from protein surfaces, organic solvents (e.g., acetone, chloroform/methanol) induce denaturation and strong non-covalent interactions between proteins, thereby driving precipitation through aggregation. Contaminants remain in solution for removal. However, protein precipitation has historically been associated with extended incubation steps, incomplete protein capture, and chemical modification of proteins and/or peptides (9, 13–16). Nevertheless, more recent methods such as S-Trap™, ProTrap XG, and SPEED (‘sample preparation by easy extraction and digestion’) have all demonstrated that combining solvent-based precipitation with filter-trapping provides a rapid means of protein capture and clean-up (17–19). Although not often employed by the aforementioned methods, the organic solvent acetonitrile (ACN) has been shown to outperform acetone in peptidomics and metabolomics studies (20–23), where rapid and efficient protein depletion is essential for analyses—often termed a protein ‘crash’ (24–27). ACN is also highly compatible with protein preparation, digestion, and LC-MS, and is widely used in proteomics research. Despite these advantages, the use of ACN protein precipitation as a tool for proteomics sample clean-up has not been fully explored.

An increasingly adopted method, SP3 (single-pot, solid-phase-enhanced sample preparation), employs magnetic beads, organic solvent denaturation, and a single reaction vessel to aggregate protein for magnet-based capture and washing (7, 28–31). The proposed mechanism of action is a hydrophilic interaction chromatography (HILIC)-like solid-phase interaction between carboxylate beads and proteins, allowing even high concentrations of detergents and salts to be removed by brief wash steps. Taking approximately 30 minutes of processing, SP3 isolates protein with a high recovery in a relatively streamlined protocol, representing a gold standard for proteomics sample preparation. The approach is parallelisable, automation-compatible, and demonstrated to be effective over a range of initial protein material inputs from 1–300 micrograms (7, 32). SP3 has proven successful in applications including phosphoproteomics, interactomics, paleoproteomics, FFPE (formalin-fixed, paraffin-embedded) and fresh-tissue proteomics, secretomics, automated proteomics, top-down proteomics, and on-bead peptide fractionation (7, 32–38). Improvements on the initially proposed method include neutral pH, solvent adjustments, and a more rapid workflow taking around 90 minutes from cells to peptides (7, 31, 32).

The main limitation to SP3 recovery is the potential for losses if protein material doesn’t completely aggregate onto magnetic beads, if aggregates are disrupted during wash steps, or if technical steps are not followed carefully (30). Losses were counter-intuitively observed at higher inputs of protein material and resulted in lower protein recovery (7, 32). These losses risk introducing variability and biasing the type of proteins captured. Large-scale protein preparations, such as those required for post-translational modification enrichment, also present a substantial cost. Further complications include the damage risk posed by stray magnetic beads carried into chromatography systems, reduced peptide yields due to binding of carboxylate beads to protease and phosphatase inhibitors in automated workflows (35), and other unwanted interactions of bead chemistries with certain chemical additives (39).

A recent finding by Batth *et al*. (34) that bead surface chemistry does not impact SP3 protein recovery suggests that HILIC-like interactions are not required to create solid-phase bead:protein interactions. Describing the mechanism as protein aggregation capture (PAC), their work evaluated several bead chemistries that gave consistent results—indicating that protein aggregation was driven by organic solvent-induced denaturation. Similarly, Lewin *et al*. (39) describe an absorption-based protocol (ABP), in which inactivation of bead surface chemistry improves protein yield by SP3, with these non-functionalised silica magnetic beads effectively capturing aggregated protein.

In this study, we build upon these findings and omit magnetic beads entirely, employing centrifugation instead of magnets to capture ACN-precipitated protein from mammalian cell lysates. We name the resulting optimised method SP4, or <u>S</u>olvent <u>P</u>recipitation <u>SP3</u>. SP4 was compared with SP3, both in the absence of beads (bead-free, BF), and with inert ∼10 µm glass beads (GB), to evaluate the effect of surface area on PAC/ABP independent of bead chemistry. Both centrifugation-based capture methods matched or outperformed SP3 in proteome coverage across a broad range of protein inputs, with glass beads generally promoting higher and more reproducible protein recovery. We provide further evidence that protein precipitation is the primary mechanism of SP3 action, and that magnetic beads, while advantageous in some settings, are generally superfluous and increase the risk of losses—especially of low-solubility (e.g., membrane) proteins and at higher protein concentrations. SP4 offers a low-cost alternative to SP3 requiring no specialised equipment or reagents, can be simplified by the omission of beads, and improves recovery of intrinsic membrane and other hydrophobic proteins, while retaining the speed and universality of SP3.

## Methods

### Material

Dulbecco’s Modified Eagle Medium (DMEM), 100x pen-strep, 100x L-glutamine, 100x MEM non-essential amino acids (NEAA), and UltraPure™ Tris were purchased from Invitrogen; HyClone™ fetal bovine serum (FBS) from Fisher Scientific; cOmplete™ mini EDTA-free protease inhibitors from Roche; BCA and peptide quantitation assays, TMTsixplex™, and LC-MS grade ACN from Thermo Scientific; HEPES from Melford Laboratories; NP-40 from Biovision; NaCl and urea from VWR International; glycerol, NaOH and LC-grade ACN from Fisher; SpeedBeads™ magnetic carboxylate modified particles 45152105050250 and 65152105050250 from Cytiva (GE Healthcare); Protein LoBind^®^ tubes from Eppendorf; Trypsin (V5111) from Promega. All other reagents were purchased from Sigma-Aldrich. Additional reagents used for validation are described in the Supplementary Methods.

### Cell culture and lysis

HEK293 cells were grown in DMEM supplemented with 10% fetal bovine serum with 1x penstrep, L-glutamine, and NEAA, in a humidified incubator set at 37 °C and 5% CO_2_. Cells were grown to 70–80% confluency and were washed twice with phosphate buffered saline. Cells were harvested by scraping, pelleted at 300 *g* and snap-frozen in liquid nitrogen.

Detergent-based lysis was performed resuspending snap-frozen cell pellets in ‘SP3 lysis buffer’ (50mM HEPES pH 8.0, 1% SDS, 1% Triton X-100, 1% NP-40, 1% Tween 20, 1% sodium deoxycholate, 50mM NaCl, 10mM DTT, 5mM EDTA, 1% (w/v) glycerol, 1x cOmplete™ protease inhibitor, 40mM 2-chloroacetamide (CAA)). Lysis was conducted by trituration with a 23-gauge needle, incubated at 95 °C for 5 min, cooled to RT for 10 min, sonicated on ice for 12x 5 s bursts with 5 s intervals and cleared at 16,000 *g* for 10 min at 4 °C. Protein concentration was estimated by BCA assay (Thermo Scientific) or NanoDrop™ 2000 (Thermo Scientific) at 280 nm, and adjusted to 5 µg/µL.

Urea-based lysis was performed in-flask with urea buffer (8M Urea, 50mM Tris-HCl pH 8.0, 75mM NaCl, 1mM EDTA, 1x cOmplete™ protease inhibitor) and lysate was snap-frozen in liquid nitrogen. Thawed lysate was triturated 20x on ice and cleared at 16,000 *g* for 10 min at 4 °C. Protein concentration was estimated by BCA assay according to manufacturer’s instructions and adjusted to 5 mg/mL. Proteins were reduced using 5mM DTT for 45 min at 25 °C and alkylated with 10mM CAA for 45 min at 25 °C.

‘SPEED’ lysis was performed as described previously (19). Briefly, an aliquoted cell pellet was lysed in 100% TFA, neutralised in 2M Tris, and reduced and alkylated with 10mM DTT and 40mM CAA. The lysate was diluted 1:1 with water and precipitated according the SP4 protocol below with 10:1 glass bead:protein ratio.

### Bead preparation

SpeedBeads™ magnetic carboxylate-modified particles (catalogue no. 45152105050250 and 65152105050250) were mixed 1:1, washed 3x using Milli-Q^®^ water, and resuspended at 50 mg/mL. Silica beads/glass spheres (9–13 µm mean particle diameter—catalogue no. 440345) were suspended at an initial concentration of 100 mg/mL in Milli-Q^®^ water, and washed 1x with 100% ACN, 1x with 100mM ammonium bicarbonate (ABC), and 2x with water. With each wash, the beads were pelleted by centrifugation at 16,000 *g* for 1 min and the supernatant discarded. Of note: approximately 50% of the beads were buoyant and did not pellet, and were carefully removed over the course of these wash steps. Metal filings were noted to be a contaminant in the beads and could be removed by magnet or acid wash, but did not impact any analyses. The beads were then resuspended in the initial suspension volume at 50 mg/mL (given ∼50% were retained) in Milli-Q^®^ water.

### SP3 or SP4 protein precipitation

Lysates were aliquoted into Protein LoBind^®^ tubes for each method and replicate. For SP4, 0.5mL tubes were used (where volumes allowed) to give the densest pellet. Either 10:1 bead:protein ratio, or the equivalent volume of Milli-Q^®^ water (for the bead-free experiments, to maintain consistent concentrations), was added to lysates and gently vortex-mixed (< 500 rpm). Samples were handled such that liquid volume was minimised. 100% ACN was added (without pipette mixing) to a final concentration of 80 % and tubes gently vortexed for 5 s. SP3 samples were incubated at 25 °C for 5 min at 800 rpm on a Thermomixer^®^ Comfort and placed on a magnetic rack for 2 min. SP4 samples were centrifuged for 5 min at 16,000 *g*. Supernatants were removed carefully, using the tube hinge to orientate pellets. Three wash steps were performed with 180 µL (or scaled equivalent) of 80% ethanol, with buffer added slowly and avoiding disturbing the beads:pellet. Each wash used either a 2 min magnetic separation (SP3) or 2 min centrifugation at 16,000 *g* (SP4).

### Proteolysis and peptide isolation

After the final wash, remaining supernatant (bar < 5 µL) was carefully removed, the bead-bound proteins resuspended by gently vortexing the beads in 100mM ABC with 1:100 trypsin:protein ratio, and placed in a sonicator bath for 5 min. For the 500 and 5000 µg samples, 1:100 TrypZean^®^ (T3568, Sigma-Aldrich) was used in place of trypsin. Samples were incubated for 18 h at 37 °C at 1000 rpm on a Thermomixer^®^ Comfort. Peptide-containing supernatants were isolated by removal of magnetic beads (magrack, SP3) or beads and insoluble debris (16,000 *g*, SP4) for 2 min.

### Peptide quantification assay

Peptide yields for optimisation were determined using the Pierce™ Quantitative Fluorometric Peptide Assay (Thermo Scientific) according to manufacturer’s instructions. For initial optimisation, samples were prepared as above, varying ACN concentration, bead:protein ratio, and centrifugation time while otherwise using 80% ACN, 5/2 min capture/wash centrifugation, and a 10:1 bead:protein ratio. Samples for each condition (n = 4) were digested in 50 µL 20mM ABC and 10 µL analysed in triplicate. For evaluation of SP3 and SP4 protein input concentrations on peptide recovery, 50 µg of HEK293 protein was aliquoted (n = 3), diluted from 5 to 0.63 µg/µL, and processed as described above.

### TMT labelling

For the 100 µg SP3 *vs*. SP4 TMT experiment, 50 µL of 100mM triethylammonium bicarbonate (TEAB) was used in place of ABC and both trypsin and Lys-C were added to a 1:100 enzyme:protein ratio. The resulting peptides were isolated by magnet or centrifugation and reaction vessels washed with 50 µL of 100mM TEAB. 0.2 mg of TMT labelling reagent was added to each sample and incubated for 1 h at RT, and treated with 8 µL of 5% hydroxylamine for 15 min at RT. Labelled peptides were vacuum-concentrated, reconstituted, and pooled.

### Peptide pre-fractionation

TMT-labelled peptides were reconstituted in 80 µL 3% v/v ACN + 0.1% v/v ammonium hydroxide and resolved using high-pH RP C18 chromatography (XBridge BEH 150mm × 3mm ID x 3.5 µm particle, Waters, Milford, MA) at 0.3 mL/min with a Dionex UltiMate™ 3000 HPLC system (Thermo Scientific) at 30 °C. Mobile phases A (2% ACN + 0.1% ammonium hydroxide) and B (98% ACN + 0.1% ammonium hydroxide) were used for a gradient of: 0–20 min (3% B), 75 min (30% B), 105 min (85% B). 70 fractions were collected in a peak-dependent manner and individually lyophilized. Fractions at the extremes of the chromatogram were subjected to solid-phase extraction (SPE) and orthogonally concatenated, giving 62 fractions for analysis.

### LC-MS analysis

Label-free analyses of SP3, BF and GB peptides were acquired using a Q Exactive™ Plus Orbitrap™ MS (Thermo Scientific) coupled with a Dionex UltiMate™ 3000 nanoHPLC system (Thermo Scientific). Peptides were separated on a reversed-phase nanoLC column (150 × 0.075 mm; Reprosil-Pur C18AQ, Dr Maisch). For each analysis the equivalent of 100 ng peptides (as a proportion of protein input) were separated using a 120 min gradient of 5–35% ACN in 0.1% FA with a flow rate of ∼300 nL/min.

Mass spectra were acquired with the following parameters for MS^1^: resolution 70,000, scan range 350–1,800 m/z, automatic gain control (AGC) target 3×10^6^, and maximum injection time 50 ms. MS^2^ spectra for 2+ to 4+ charged species were acquired using: HCD fragmentation, top 10, resolution 17,500, AGC 5×10^4^, maximum injection time 100 ms, isolation window 1.2 m/z, and normalized collision energy (NCE) of 27. The minimum AGC target was set at 2×10^3^, which corresponds to a 2×10^4^ intensity threshold.

TMT-labelled high pH peptide fractions were analysed by Orbitrap™ Eclipse MS (Thermo Scientific) with on-line separation on a reversed-phase nanoLC column (450mm × 0.075mm ID) packed with ReprosilPur C18AQ (Dr Maisch, 3 µm particles) at 40 °C. A 60 min gradient of 3–40% ACN, 0.1% FA at 300 nL/min was delivered via a Dionex UltiMate™ 3000 nanoHPLC system. Mass spectra were acquired in SPS MS^3^ mode using a 3 s cycle time with the following settings: MS^1^—120k resolution, max IT 50 ms, AGC target 400,000; MS^2^—IW 0.7 CID fragmentation, CE 35 %, max IT 35 ms, turbo scan rate, AGC target 10,000; MS^3^— HCD fragmentation, CE 55 %, 30k resolution, max IT 54 ms, AGC target 250,000.

### Data analysis

LC-MS raw files were processed with Proteome Discoverer™ 2.5 using Sequest HT and Percolator, searching against UniProt Human Swissprot (UniProtKB 2021_01) and a PD contaminant list (2015_5). Default settings were used, allowing two missed tryptic cleavages, with carbamidomethyl (C, fixed), oxidation (M, variable), acetyl (protein N-term, variable), and, for the isobaric-labelled experiment, TMT 6-plex (K, peptide N-term, fixed). FTMS and ITMS spectra were searched with 0.02 and 0.5 Da fragment mass tolerances, respectively. Proteome Discoverer™ was used to determine protein and peptide identifications (q < 0.01), CV values, TMT quantification, and differential abundance p-values (t-test). Minora feature detector was used for label-free quantification. No normalisation was applied to assess technical effects. Additional t-tests were performed in Microsoft Excel with no assumption of equal variance. *R*^2^ values were determined as the squared Pearson product-moment correlation coefficient, using Microsoft Excel. The MS proteomics data have been deposited to the ProteomeXchange Consortium (http://proteomecentral.proteomexchange.org) via the PRIDE partner repository (40) with the dataset identifier PXD028732. Proteomics data are detailed in Table S1. Gene Ontology term enrichment analysis and functional annotation enrichment was performed with DAVID version 6.8 using a background of both the full human proteome and all TMT-study identified proteins. Terms were filtered to include those with Benjamini-adjusted significance (p < 0.05). For protein solubility analysis, the UniProt Human Swissprot proteome was submitted to the CamSol Intrinsic tool for the calculation (at pH 7) of protein solubility and generic aggregation propensity, with a score generated for each protein sequence (41). Hydrophobicity (GRAVY score) was calculated by the PROMPT tool (42), and isoelectric points from ProteomePI (43).

### Manuscript preparation

This manuscript was prepared in Overleaf (http://www.overleaf.com), using the [HenriquesLab bioR*χ*iv template](https://www.overleaf.com/latex/templates/henriqueslab-biorxiv-template/nyprsybwffws#.Wp8hF1Cnx-E).

## Results

### Single-pot solvent precipitation with acetonitrile provides effective protein capture and clean-up

Building on previous mechanistic observations of SP3, we wanted to explore further the hypothesis that protein capture observed in SP3 is a product of solvent-induced denaturation and aggregation, rather than being dependent on bead surface chemistry (34, 39). These findings led to the suggestion that the aggregation observed in SP3 may be identical in mechanism to that of solvent-induced protein precipitation. Given the risks of losses presented by reliance on protein–bead adhesion, and the effective use of ACN to precipitate out proteins in peptidomics and metabolomics, we considered that centrifugation to capture ACN-precipitated protein may provide an effective means of sample clean-up for proteomics. The SP3 method was therefore adapted for direct comparison of centrifugation *vs*. magnetic protein aggregation capture (Fig. 1).

**Fig. 1.**
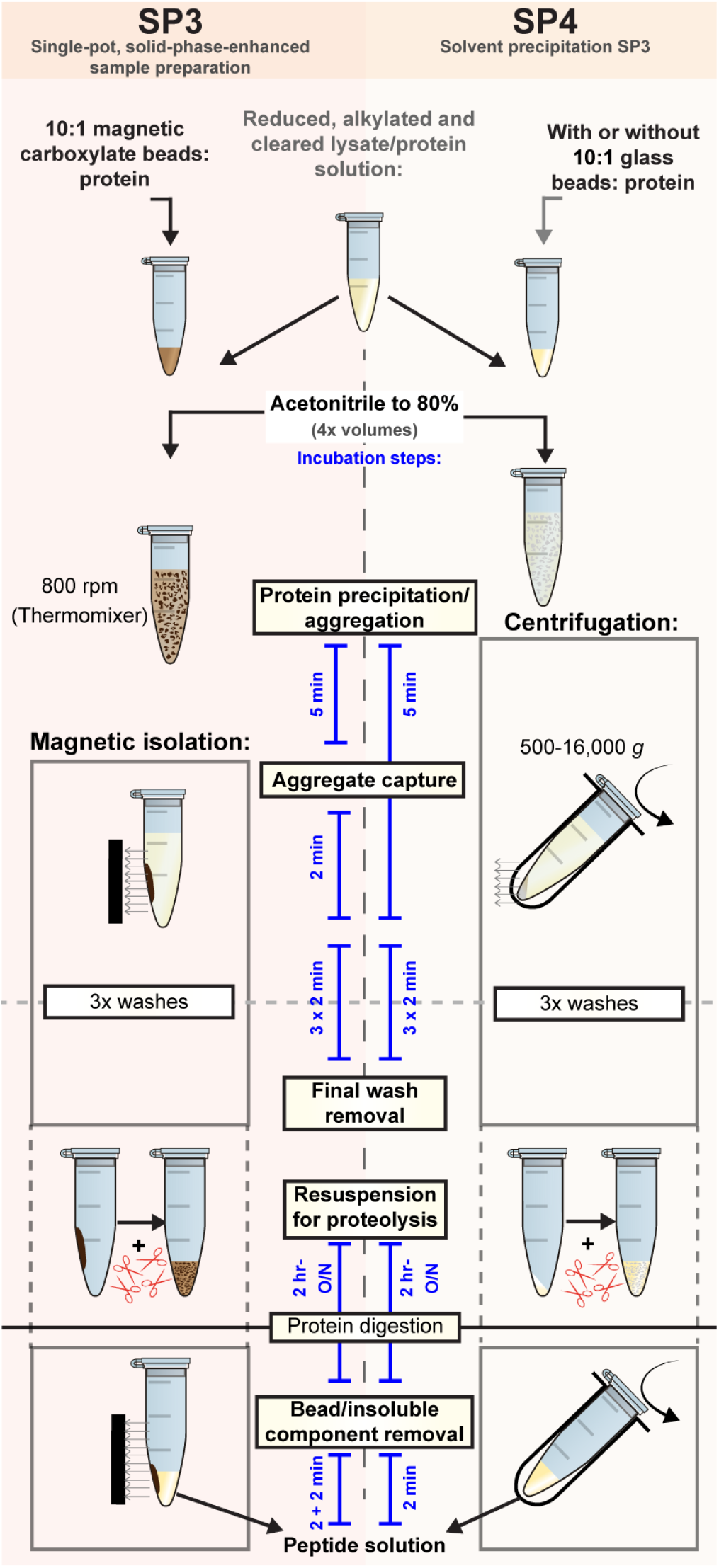
SP3 and SP4 workflows. For both approaches, a protein solution is adjusted to 80% acetonitrile to trigger protein denaturation and aggregation. SP3 captures aggregates with carboxylate magnetic beads (10:1 beads:protein), whereas SP4 uses centrifugation and, optionally, 10:1 glass beads:protein. Captured aggregates are washed 3x without resuspension, and recaptured by magnet or centrifuge for 2 min. The washed protein aggregates are then resuspended for proteolysis, and the peptide supernatant cleared by magnet and/or centrifugation isolation of the insoluble material and beads. These peptides are suitable for downstream applications such as direct LC-MS, PTM-enrichment, fractionation and/or isobaric labelling.

We named this SP4, for *<u>S</u>*olvent *<u>P</u>*recipitation *<u>SP3</u>*. Two variants were devised: one without any beads (bead-free, BF), thus relying on precipitation alone, and a second with inert, low-cost, silica particles (hereafter termed glass beads, GB), allowing us to explore the role of surface area independently of bead chemistry. Importantly, for the purposes of direct comparison to SP3, SP4 used a single-pot reaction, 5- and 2-minute protein capture steps and, for the glass beads, a 10:1 bead:protein ratio. The protocol was also adapted to incorporate many of the recent optimisations to SP3, including neutral pH, 80% ACN for aggregation, and no disturbance of the protein–bead aggregates (7, 31, 32).

A broad range of SP4 parameters were evaluated by peptide yield, including 40–95% ACN, 0:1 to 160:1 glass bead:protein, and 30 seconds to 20 minutes centrifugation time (Fig. S1). These initial experiments demonstrated that 80% ACN, a 10:1 glass bead to protein ratio, and 5- and 2-minute protein capture steps did indeed offer the most effective protocol. Optimal conditions were less pronounced when these parameter ranges were measured by proteomics (n = 1), but broadly confirmed that the parameters were highly effective for those conditions equivalent to SP3. Notably, these optimisations additionally highlighted that: (1) the use of glass beads improved protein yield; (2), precipitation and centrifugation times as low as 30 seconds still provided > 90% of the maximum peptide yield; and (3) there was no observed interference from any detergent contaminants for any conditions tested. Rapid protein aggregate capture by centrifugation-based SP4, using parameters equivalent to SP3, therefore provided a viable option for the preparation of samples for proteomics analysis.

### Centrifugation outperforms magnetic capture of solvent-induced protein aggregates

To understand differences between solvent-induced protein precipitation and aggregation capture by magnetic carboxylate beads, cell lysate was processed by bead-free and glass bead SP4 alongside SP3 across a broad range of protein inputs, from 1–5000 µg (Fig. 2A & S2; Table S1). Overall, SP4 consistently provided equivalent or higher-quality proteome data than SP3. The inclusion of glass beads during aggregation provided additional improvements to protein identifications and reproducibility in most instances. Median protein *R*^2^ values were 0.970, 0.980, and 0.993 for SP3, BF, and GB, respectively (Fig. S3). For both bead-free and glass bead SP4 methods, more proteins were recovered at our significance threshold (Fold-Change (FC) > 2 and adjusted p < 0.05) *vs*. SP3 (Fig. 2B & S4).

**Fig. 2.**
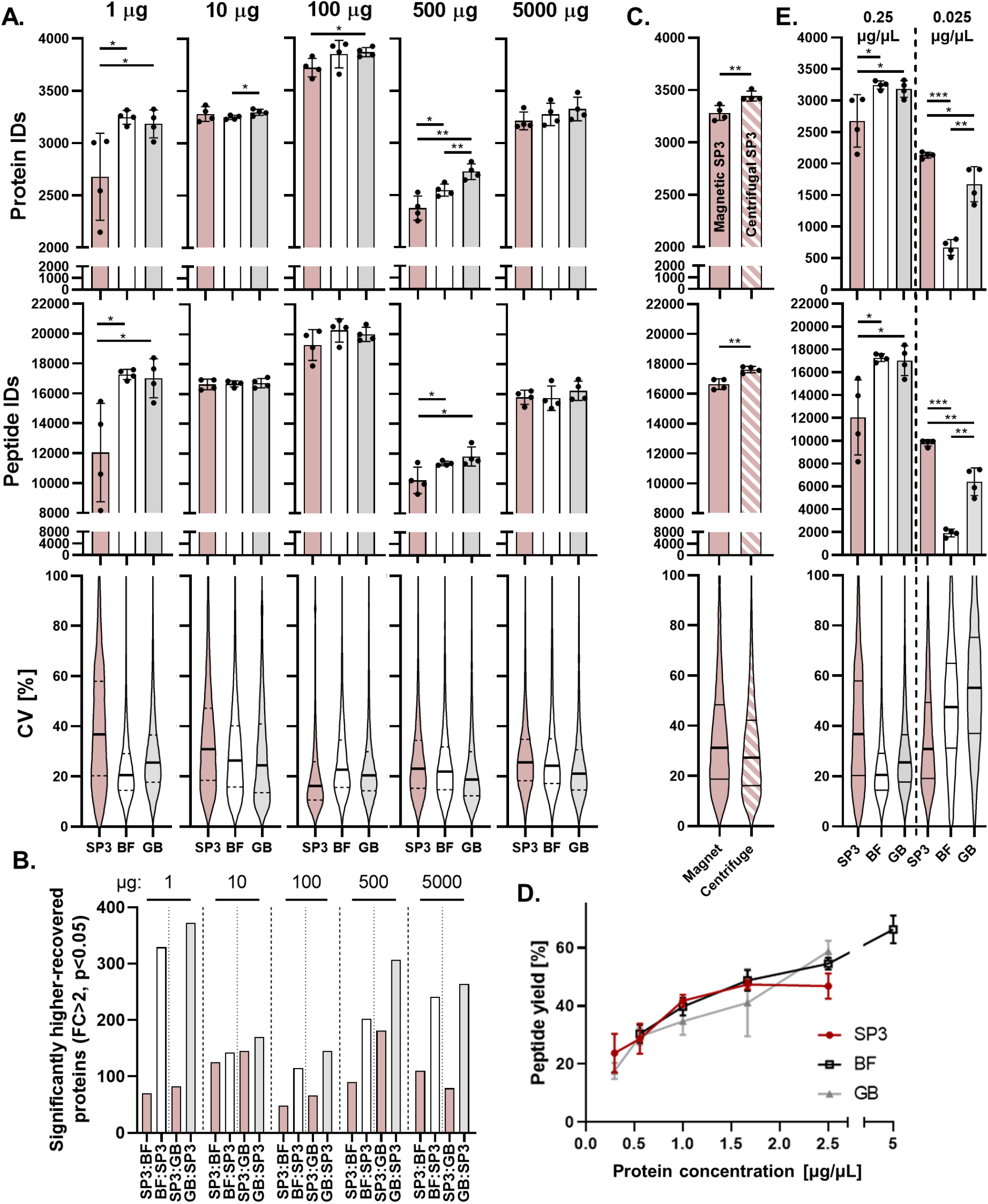
Comparison of magnetic protein–bead aggregate capture (SP3) with centrifugation of aggregates formed by solvent precipitation SP3 (SP4) in the absence (bead free, BF) and presence of inert 10 µm glass beads (GB) at a 10:1 bead:protein ratio. **A**. Protein and peptide identifications and peptide coefficient of variance (CV) for SP3, BF and GB preparations (n = 4) ranging from 1–5000 µg using SP3 and solvent precipitation SP3 (SP4). Samples were processed at a final concentration of 0.25 µg/µL (1–10 µg) and 2.5 µg/µL (100–5000 µg). 500 and 5000 µg samples were digested with TrypZean™. **B**. The numbers of proteins found to be differentially recovered (FC > 2 and p < 0.05) by each method, for the experimental sets in A. **C**. 10 µg of protein was processed by SP3 (n = 4) with the protein–bead aggregates captured by centrifugation at 16,000 *g*, compared with standard magnetic capture. **D**. 50 µg of protein was processed by SP3, BF and GB methods across a range of protein concentrations, representative of that of the final volume, including the volume from the bead suspension for SP3 and GB. The resulting peptides were measured by peptide assay. It was possible to evaluate the bead-free method at a 5 µg/µL protein concentration, as no volume from a bead suspension needed to be added. **E**. 1 µg preparations were processed at a concentration of 0.025 µg/µL compared with 0.25 µg/µL and evaluated by proteomics. Significance was measured by t-test; * p < 0.05, ** p < 0.01, *** p < 0.001

Our lowest protein input (1 µg) highlighted variable protein losses in SP3, with SP4 providing significantly greater coverage and lower coefficients of variation (CV) (p < 0.05). For 10 µg preparations, no method offered a clear advantage; however, the inclusion of glass beads provided significantly more protein IDs (p < 0.05) and greater reproducibility *vs*. bead-free. Although more proteins were identified for SP4 (p < 0.05 for GB), for the 100 µg preparations, SP3 did offer lower CV values. These SP4 variances were still below all other variances measured for SP3. The 500 µg preparations demonstrated a significant advantage of SP4 over SP3 (p < 0.05), and of glass beads over bead-free (p < 0.01). 5000 µg preparations demonstrated a similar trend, but without significance. Note that 500 µg and 5000 µg preparations—both digested with TrypZean™ instead of proteomics-grade trypsin—demonstrated a marked drop-off in total protein identifications *vs*. lower inputs, consistent with previous SP3 studies (7, 32). Missed cleavages were also generally reduced in SP4 preparations, particularly in the presence of glass beads (Fig. S2).

Next, we combined magnetic bead-based SP3 with centrifugation (Fig. 2C & S5) to demonstrate that centrifugation outperforms magnetic capture of protein–bead aggregates, with significantly increased protein and peptide identifications (p < 0.01) and lower variability.

An important observation was the effect of protein concentration, and therefore total aggregation reaction volume, on protein recovery. Fig. 2D demonstrates the losses of recovered peptide material resulting from lowering protein concentration during aggregation. Each 2-fold dilution resulted in, on average, a 15% and 20% loss of material for SP3 and SP4, respectively. Notably, SP4 outperformed SP3 at higher concentrations, while SP3 marginally outperformed SP4 at some lower concentrations. This effect was evaluated by proteomics with a 1 µg input prepared at a concentration of 25 ng/µL, compared with the previous 250 ng/µL preparation (Fig. 2E & S5A). At this reduced protein concentration, SP3 outperformed SP4, identifying over three times as many proteins observed in the absence of beads—highlighting a noteworthy limitation of the SP4 method.

Several additional aspects of the SP4 protocol were evaluated to identify and understand potential improvements or adaptations. First, given the similarity of SP4 to acetone precipitation, SP3 and SP4 preparations were performed using 80% ACN alongside 80% acetone, and otherwise identical conditions (Fig. S5B). No significant differences were observed between the two solvents. Next, the use of lower SP4 centrifugation speeds—more readily compatible with larger volumes and high-throughput plate-based preparations—were considered (Fig. S5D). 500 *g* was not only effective, but provided improvements to the peptide yield in both the presence and absence of glass beads, suggesting less-dense and therefore more trypsin-porous pellets. Finally, we assessed and confirmed the compatibility of SP4 with detergent-free lysis methods including lysis by trifluoracetic acid using the recently described ‘SPEED’ protocol (Fig. S5E), as well as in buffers containing the common proteomic solubilising agent urea (Fig. S5F). Notably, SPEED combined with SP4 provided greater proteome coverage and lower variability than the equivalent glass bead controls using detergent-based buffer.

Together, these findings suggest that centrifugation-based protein aggregate capture by SP4 robustly offers advantages over the dependence on magnetic bead–aggregate interactions of SP3 (except in circumstances where protein concentration is very low), and confirm the compatibility of a broad range of cell lysis and aggregate-capture parameters.

### Deep proteome profiling identifies superior recovery of membrane and low-solubility proteins by SP4

Having established that SP4 provided yields equivalent or superior to SP3 amongst the 3000 most abundant proteins, we designed an isobaric labelling study to provide a deeper, more global, proteome coverage comparing protein capture by SP3 and SP4 (Fig. 3A). 100 µg of peptides were prepared from a single starting lysate by SP3, bead-free, and glass bead solvent precipitation in duplicate, labelled with TMTsixplex™, and analysed by 2D LC-MS^2^ using synchronous precursor selection (SPS) and MS^3^ quantification of 62 fractions. In total, this experiment quantified 9,076 proteins across all six samples.

**Fig. 3.**
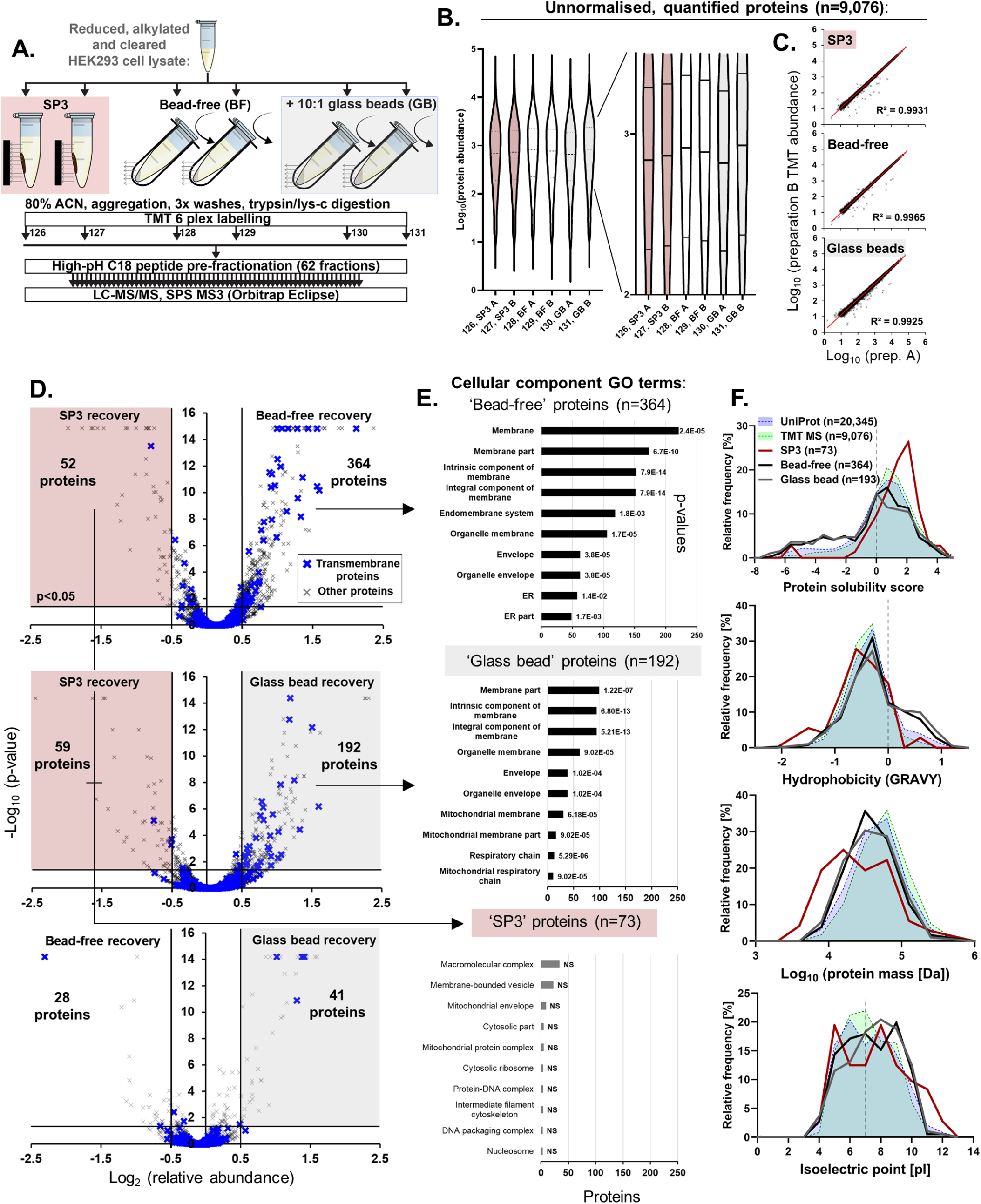
Deep proteome profiling comparing SP3 and SP4 (with and without 10 µm glass beads) by isobaric labelling. **A**. Experimental workflow applied to investigate the differences between SP3 and SP4 to a depth of 9,076 proteins. **B**. Proteome-wide relative protein abundances observed for the six sample preparations, inset; zoomed to highlight the differences in median and quartile values. **C**. Correlation between protein abundances for sample preparation method replicates. **D**. Protein recovery observed to be more effective by each of the preparation approaches. **E**. Gene Ontology term enrichment analysis of those proteins more effectively isolated by each method detailing the number of proteins matching each term and the Benjamini-corrected enrichment p-value (NS = not significant). **F**. The relative frequency distributions of protein properties amongst those proteins significantly enriched by BF, GB or SP3 preparations. Both the human UniProt Swissprot proteome and the TMT quantified proteome are displayed as backgrounds.

Protein recovery was higher in both centrifugation-based SP4 preparations relative to SP3, as measured by TMT. Although glass bead-based recovery was higher on average, it was the more variable of the three approaches (Fig. 3B). Comparing protein abundance between replicates (Fig. 3C & S6) demonstrated a high degree of correlation for all three methods, with bead-free having the greatest reproducibility (*R*^2^ = 0.9965), followed by SP3 (*R*^2^ = 0.9931) and glass beads (*R*^2^ = 0.9925). Assessing differentially recovered proteins by volcano plot (Fig. 3D) further demonstrated a greater overall protein yield by SP4 and a larger number of proteins more significantly recovered for both bead-free (364) and glass bead (193) approaches. Only 73 proteins had greater recovery by SP3 versus both SP4 variants, and very little differential recovery was observed between SP4 variants. Functional annotation enrichment analysis of these three protein subsets (Fig. 3E & S6) identified a strong trend of SP4 differentially-recovered proteins annotated with terms such as ‘membrane’ (n = 221/364, p = 2.4×10^−5^) and ‘intrinsic component of membrane’ (n = 153/364, p = 7.9×10^−14^) for both variants. Across the SP4 preparations, approximately half of all proteins with additional recovery were annotated to be membrane-associated—twice as many as would be expected by chance (Fig. 3D, blue crosses). Of these SP4 differentially-recovered proteins, 111 (28 %) were multi-pass membrane proteins, with 953 (9.2 %) identified in the TMT proteome relative to a 14% frequency in the human proteome. No terms were significantly enriched for the SP3 differentially-recovered proteins. The physicochemical property distributions of these SP3 and SP4 differentially-recovered protein subsets were also evaluated relative to the TMT and human UniProt proteomes (Fig. 3F), additionally highlighting a significant enrichment by SP4 of hydrophobic proteins and those with a lower predicted solubility (p < 0.0001). SP3, although limited by low protein numbers for this analysis, did suggest some enrichment of high-solubility, hydrophilic, and smaller proteins. Taken together, our analysis suggests that centrifugation-based SP4 provides superior recovery to magnet-based SP3 across the whole proteome, and that membrane and other low-solubility proteins appear more effectively captured by centrifugation-based approaches.

### SP4 matches or outperforms SP3 independent of user

To confirm that SP4 was not dependent on any single user or setting, the protocol was shared with three collaborators to compare with SP3, two of whom were regular users of SP3 (Fig. 4). Lab A first found that glass bead SP4 outperformed SP3 with two different carboxylate-modified magnetic particles (ReSYN or SpeedBeads™) at lower protein inputs, with roughly equivalent performance at higher inputs. They also compared glass bead SP4 with overnight precipitation using acetone, demonstrating similar performance. Next, they compared trypsin with and without Lys-C for acetone precipitation and SP4. Lys-C addition substantially reduced variance in both approaches, with SP4 consistently providing the highest number of protein and peptide IDs (p < 0.01). Lab B, performing SP3 for the first time, prepared 25 µg in triplicate and found the approaches roughly equivalent. Lab C processed two independent n = 5 comparisons of SP3 and glass bead SP4 using 50 µg of E14 murine embryonic stem cell lysate. For both experiments, approximately 100 more proteins were identified by SP4 (p < 0.001), even though the number of peptides did not significantly differ between comparisons. The variances were marginally lower for SP4 in both comparisons. These validations demonstrate that advantages seen with SP4 vs. SP3 are achievable by multiple users and settings, highlighting that the protocol provided is both robust and straightforward to adopt.

**Fig. 4.**
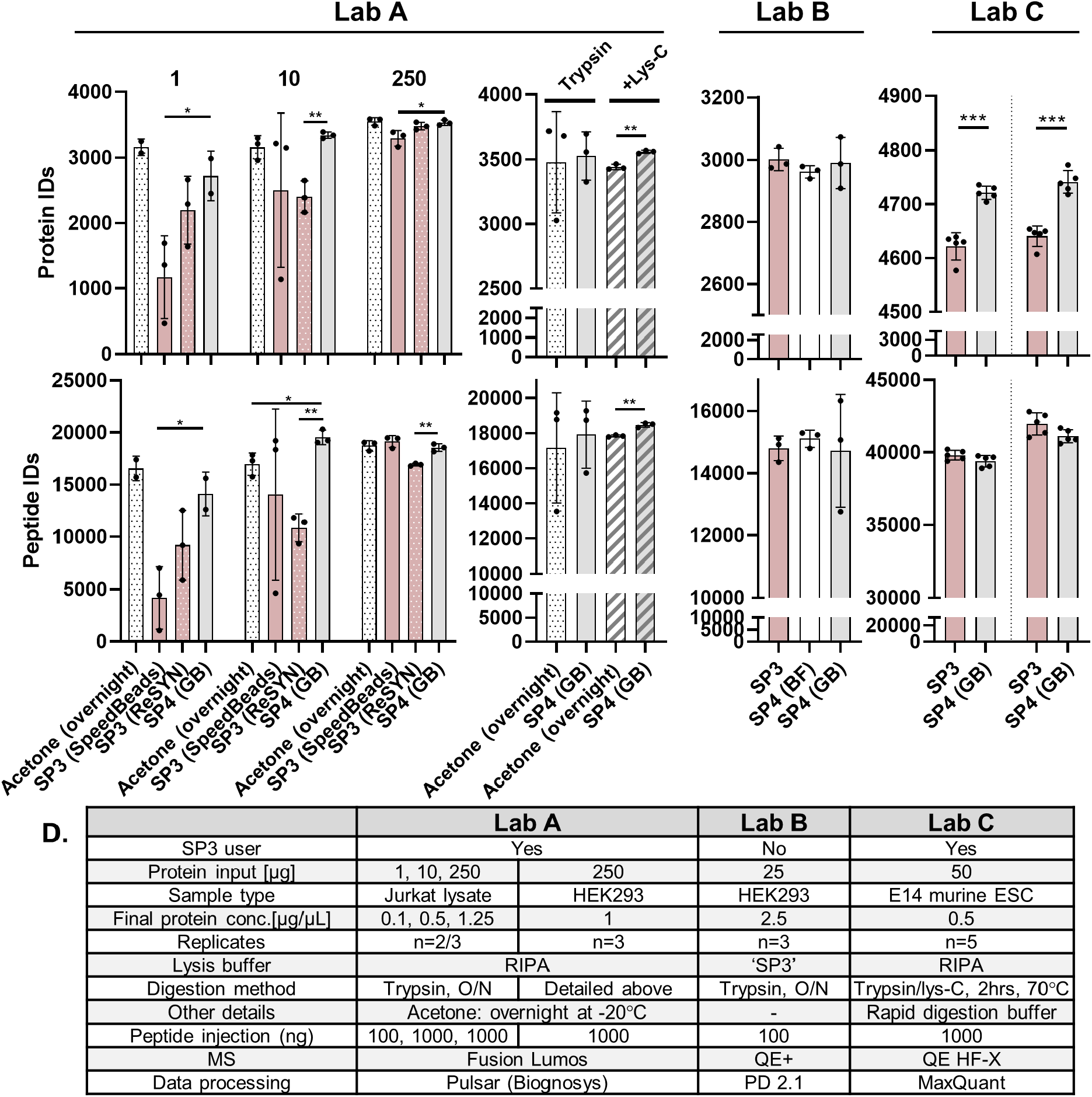
Independent validation of SP4 protein clean-up. The SP4 protocol was provided to three collaborators, with each comparing variants of SP4 to SP3. **A**. Lab A performed SP3 with either SpeedBeads™ or ReSYN carboxylate magnetic beads compared with overnight acetone precipitation and SP4 with glass beads (GB) for 1, 10 and 250 µg preparations. Additionally, acetone precipitation and SP4 were compared for trypsin in the presence and absence of Lys-C. **B**. Lab B processed 25 µg of HEK293 lysate for SP3, bead-free (BF) and GB protocols. **C**. Lab C processed two independent n = 5 comparisons of SP3 and glass bead SP4 using 50 µg of E14 murine embryonic stem cell lysate. **D**. Summary of the methodologies used by each lab. Significance was measured by t-test; * p < 0.05, ** p < 0.01, *** p < 0.001

## Discussion

SP3 is one of the most effective means of proteomics sample capture and clean-up currently available. However, the reliance on stable aggregation of proteins onto magnetic beads remains a source of variability and loss. This work demonstrates that magnetic protein aggregate capture by SP3 and solvent-induced protein precipitation by SP4 perform very similarly, suggesting precipitation as the primary mechanism of SP3. Protein aggregate capture by centrifugation broadly improved proteome quality and reproducibility independent of protein input (Fig. 2, 3, & 4), particularly amongst low-solubility and membrane proteins (Fig. 3)—further highlighting general and specific losses resulting from magnetic bead-based capture in SP3. SP4 was effective in a range of lysis buffers with high concentrations of detergents, neutralised acid, and urea (Fig. 2, 4, & S5E-F), exhibiting no contamination carryover. SP4 was also effective for deep proteome profiling (Fig. 3), and was successful in the hands of multiple users with varying levels of SP3 experience (Fig. 4).

Previous results (34, 39), and those presented here, demonstrate that specific bead surface chemistry was not required for protein aggregate capture. Indeed, even in the absence of beads, protein precipitation consistently captured at least equivalent levels of material to that of SP3 beads (Fig. 2, 3, & 4). Importantly, SP3 did appear advantageous at low protein concentrations, with carboxylate beads apparently acting as a surrogate aggregation surface, thereby improving recovery (Fig. 2E). Although glass beads also provided an advantage to recovery compared with bead-free at low protein concentrations, this improvement was not quite to the same extent as SP3 beads. Therefore, surface chemistry appears to become important for dilute samples where aggregation can be accelerated by additional nucleation points—in agreement with previous observations that higher bead inputs are beneficial at lower concentrations (34). Conversely, SP4 outperformed SP3 at higher protein concentrations (Fig. 2D-E), consistent with previous observations of losses in SP3 at higher inputs (7, 32). This lower efficiency of SP3 capture at higher protein inputs may be explained by rapid auto-nucleation of protein aggregates out-competing adhesion onto the surface of carboxylate beads—creating non-magnetic particulates in suspension that are lost through washes. Notably, SP3 conducted in low-volume aggregation reactions (Fig. 2A, 1 µg) demonstrated greater variability than higher-volume reactions (Fig. 2E; also observed by Lab A’s validation in Fig. 4A, 1 µg). Our findings suggest that protein concentration and aggregation reaction volume should be carefully considered in the experimental design of future applications of SP3 and SP4.

The additional proteins recovered by SP4 (Fig. 2, 3, & 4) seem likely to arise from the centrifugal capture of fragile or less adhesive aggregate particles not efficiently captured by SP3 beads. To minimise protein shearing during wash steps for SP3 and SP4, washes were applied carefully without pellet resuspension. Therefore, improvements to proteome quality observed with SP4 suggest that some proteins and their aggregates do not adhere strongly to SP3 beads or other aggregates—demonstrated by improved recovery when centrifuging the magnetic SP3 beads (Fig. 2C). A significant enrichment of membrane-related proteins identified by deep proteome profiling for SP4 (Fig. 3) additionally suggests that hydrophobicity may give rise to increased solubility in organic solvent, and therefore resistance to aggregation. This is consistent with previous observations of hydrophobic proteins having a greater resistance to organic solvent-induced precipitation (44, 45). The effects of other parameters, such as ACN concentration, on the observed enrichment of membrane proteins for SP4 vs. SP3 warrants further investigation to understand why these proteins appear more readily lost during SP3. Given the additional challenges posed during sample handling of highly hydrophobic proteins (such as those in cell membranes, or unfolded/misfolded proteins), and their fundamental importance in drug discovery and biology, this advantage of SP4 is important to consider in experimental design.

The addition of a chemically inert surface area in the form of glass beads was considered to provide an aggregation capture surface similar to that of the absorption-based protocol onto unfunctionalized silica beads (39). This idea is also supported by tube wall ‘coating’ from surface aggregation that we observed in SP3 preparations. Improvements arising from the addition of glass beads very likely stems from this phenomenon. Following centrifugation, glass bead-protein aggregates formed a dense matrix that may be more resistant to losses from pipetting. Less protein was observed to adhere to tube walls when glass beads were present in precipitation reactions, suggesting they out-compete tube walls as nucleation surfaces, reducing this potential source of losses. Despite these advantages, some experiments demonstrated greater variability with glass beads (Fig. 3 & 4). We noted that the glass beads settled out of solution more quickly than the magnetic SP3 beads, which would potentially present a source of variability. Adding beads pre-suspended in ACN may improve dispersion uniformity and eliminate the extra dilution introduced by the bead suspension. Glass beads appeared to serve a secondary function in disrupting the density of the pellet and aggregates, thereby facilitating resuspension, providing a greater surface area accessible to trypsin, and lowering the rate of missed cleavages (Fig. S5). Additionally, the glass beads are approximately 1/1000^th^ the cost of SP3 beads, present greater chemical compatibility (e.g., with amine-reactive reagents), are not destroyed by freezing (allowing the addition of pause points in the protocol), and appeared easier to remove from samples due to their larger size. SP4 with glass beads was fully compatible with rapid digestion approaches in as little as 2 hours (Fig. 4C).

A key advantage of SP4 over SP3 is its near-limitless scalability in both volume and protein input, limited only by the size of the available reaction vessel being used; therefore, SP4 potentially provides benefits beyond micro-scale proteomics applications. The absence of beads may also lend itself to approaches where a bead-removal step is undesirable and potentially enables protein aggregation, washes, digestion, and LC-MS loading all in a single reaction vessel— making it a true ‘single pot’ method. By adopting a single-pot strategy, SP4 also avoids the need for centrifugal filters and additional sample tubes which may present surfaces for losses (e.g. FASP, S-Trap™, ProTrap XG) and their associated costs. SP4 of TFA-solubilised, Tris-neutralised proteins (SPEED method) notably provided the lowest median CV% of any of the 10 µg preparations (Fig. S5E). The Tris neutralisation step causes formation of a very fine protein precipitate, which likely acts as a nucleation surface similar to beads with the additional advantages offered by the SPEED method.

Compared with many other established protein precipitation protocols, SP4 substantially reduces sample processing time, employs a more protein-compatible solvent, and provides the option to use glass beads; offering the advantages, discussed above. The observation of no significant difference between ACN and acetone as the solvent driving aggregation and precipitation (Fig. S5B) suggests SP4 may be independent of organic solvent type. However, further investigation is required, e.g., at lower protein inputs and different solvent concentrations. Lab A found that compared with overnight acetone precipitation, a 5-minute combined precipitation and centrifugation step for SP4 provided equivalent or greater yields in most circumstances (Fig. 4A).

At very low protein concentrations, SP4 did appear more susceptible to reduced recovery. Although this is partially mitigated by glass beads, and could be improved with longer precipitation steps, SP3 beads may still be advantageous in low protein concentration applications—although our data suggest these are also similarly impacted by dilution (Fig. 2D). For low-concentration samples the use of non-magnetic carboxylate beads with centrifugation would likely provide the benefits of both approaches. Another major strength of SP3 is its use in automated workflows. For SP4, we demonstrate that low centrifugation speeds—broadly compatible with 96-well plates—are effective, potentially allowing high scalability of the approach. Initial attempts suggested that the preparation of multiple 96-well plates, from plated samples to the start of digestion, would take less than an hour.

A noteworthy limitation of SP3, SP4, and protein precipitation-based methods in general remains the loss of very low mass peptide material, potentially excluding relevant biological materials from analyses, e.g., those enriched by similar ACN-based methods for peptidomics. On the other hand, this effect may result in the exclusion of small, degraded protein fragments from peptide samples that increase sample complexity and are less relevant in the study of protein functionality.

SP4 undoubtedly has the potential for further optimisation. The precipitation step, for instance, may be enhanced by cold temperatures and longer centrifugation at slower speeds (hinted at in Fig. S5D). The trade-off between a denser aggregate pellet and the ease of re-suspension for trypsin accessibility may be worthy of further exploration (Fig. S5D), although Lys-C and rapid digestion buffers appear to be effective solutions (Fig. 4A, C). Another parameter for improvement is the balance between ACN concentration for more complete precipitation and the losses this presents from exponential reaction volume increases (Fig. 2D-E & S1). However, these would likely require deep proteome profiling to provide a wider understanding of the effects on differential protein recovery.

### Concluding Remarks

SP3 undoubtedly provides an efficient and effective means of protein capture and clean-up, although some limitations to the protocol remain, most notably the risk of variable losses during aggregation and wash steps. Building on promising developments to the SP3 protocol, we have evaluated and validated a method aimed at addressing these issues by replacing reliance on magnetic bead capture with centrifugation, thereby lowering cost, improving recovery (especially of low-solubility and membrane proteins), and increasing reproducibility. This work also builds on previous suggestions that bead chemistry is dispensable for SP3. Aggregation interactions observed between carboxylate beads and proteins resulted in highly similar proteomes to the interactions during protein-protein aggregation and precipitation alone. Inert surfaces were also capable of promoting protein aggregation capture. At low protein concentrations the availability of carboxylate surfaces in SP3 are beneficial as nucleation points, while at high protein concentrations, precipitation appears to out-compete bead surfaces, risking losses of unbound precipitate during washes. We hope that these findings will extend the options available for proteomics sample clean-up, deepen mechanistic understanding of SP3, and continue to encourage further development of universal methods for proteomics workflows.

## ACKNOWLEDGEMENTS

The authors would like to thank Richard Kay for useful discussions about peptidomics methods and John Timms, who helped to support early work that inspired this study. HEJ and RSS are funded by Institute Strategic Programme Grant BB/P013384/1 from the BBSRC.

## List of Supplementary Material

### Supplementary Methods for Fig. 4

**Figure S1**. Evaluation of a range of SP4-related variables by peptide quantification assay (n = 4) and proteomics analysis.

**Figure S2**. Additional measures of proteome quality and protein recovery for the comparison of SP3 to SP4 with (GB) and without glass beads (bead-free, BF) across a range of protein inputs (see also Fig. 2).

**Figure S3**. Protein and peptide LFQ *R*^2^ values of recovery for SP3 to SP4, summarising the data presented in Fig. 2.

**Figure S4**. Protein recovery observed to be more effective by SP4 variants versus SP3, summarised in Fig. 2.

**Figure S5**. Additional experiments exploring the mechanism and potential of bead-free (BF) and glass bead (GB) SP4.

**Figure S6**. Additional measures of quantitative proteome quality for the comparison of SP3 to SP4 with and without glass beads using TMTsixplex™ and SPS MS^3^, from Fig. 3.

**Figure S7**. DAVID-derived term enrichment and clustering for those proteins observed more significantly recovered by SP3 and SP4 by TMT quantification.

**SP4 Protocol**. Step-by-step protocol for bead-free (SP4-BF) or glass-bead (SP4-GB) sample preparation.

**Table S1**. Detailed tables and summaries of proteomics findings.

## Supplementary Methods for Fig. 4

### Lab A

#### Experiment 1

**Lysate preparation**. Jurkat cell lysate was prepared and diluted with RIPA lysis buffer to 1.25 µg/µL. Three different masses of Jurkat protein lysate in RIPA lysis buffer were prepared: 250 µg (1.25 µg/µL), 10 µg (0.5 µg/µL) and 1 µg (0.1 µg/µL). Each experiment was performed as at least three biological replicates. **Sample preparation**. Cell lysates were reduced (5mM DTT, 30 min, 25 °C) and alkylated (5mM iodoacetamide, 30 min, 25 °C in the dark). Proteins were recovered by one of four methods. For the acetone precipitation method, proteins were precipitated by adding ice-cold acetone (4x sample volume, overnight, -20 °C). Protein pellets were obtained by centrifugation (18,000 *g*, 10 min, 4 °C) and washed 2x with the same volume of ice-cold 80% acetone/water (with sonication between washes). The final wash liquid was aspirated, and samples were air-dried for 20 min. Each sample was resuspended in 50 mM HEPES (250 µL for 250 g and 20 µL for 10/1 µg). Samples were sonicated and vortexed to re-dissolve the pellet. For the Sera-Mag™ SP3 bead method, a stock of SP3 beads was prepared at 50 mg/mL by combining equivalent volumes of hydrophobic bead slurry and hydrophilic bead slurry. The resulting slurry was washed 3x with water and 3x with 50mM HEPES. SP3 beads were added to cell lysate in bead/protein ration of 10:1 (w/w) and distributed through gentle pipetting. The volume of the mixture was doubled with absolute ethanol and shaken (10 min, 800 rpm). Tubes were placed on the magnetic separator and allowed to separate. The beads with precipitated protein were washed 3x with the same volume of 70% ethanol/water and then re-distributed with 50mM HEPES to give 250 µg (1.0 µg/µL), 10 µg (0.5 µg/µL), and 1 µg (0.1 µg/µL). For the ReSYN HILIC bead method, a stock of ReSYN HILIC beads was supplied at 50 mg/mL. The resulting slurry was washed 3x with water and 3x with 50mM HEPES. ReSYN beads were added to cell lysate at 10:1 bead:protein ratio (w/w) and distributed through gentle pipetting. The volume of the mixture was doubled with 50mM HEPES/30% acetonitrile mixtures (binding buffer), and the tubes were shaken (30 min, 800 rpm). Tubes were placed on the magnetic separator and allowed to separate. The beads with precipitated protein were washed 3x with an equivalent volume of 95% acetonitrile and then re-distributed with 50mM HEPES to give 250 µg (1.0 µg/µL), 10 µg (0.5 µg/µL), and 1 µg (0.1 µg/µL). For the glass bead method, 100 mg of glass beads was distributed in 1mL of UltraPure*TM* water. This slurry was vortexed and centrifuged (16,000 *g*, 2 min, 4 °C). The buoyant beads were gently aspirated to leave a glass bead pellet. This process was repeated 1x with acetonitrile, 1x with 50mM HEPES, and 2x with UltraPure*TM* water. On the final wash, beads were resuspended in 1mL of UltraPure*TM* water, and a bead concentration of 50 mg/mL was assumed. Glass beads were added to cell lysate at 10:1 bead:protein ratio (w/w) and distributed through gentle vortexing. ACN was added to a final concentration of 80 %. Upon addition of ACN, the mixture was again gently vortexed, and the tubes were centrifuged (16,000 *g*, 3 min (2x, with tubes spun in between), 4 °C). The liquid was gently aspirated, and the beads were washed 3x with the equivalent volume of 80% ethanol/water. Beads were re-distributed with additional sonication with 50mM HEPES to give 250 µg (1.0 µg/µL), 10 µg (0.5 µg/µL), and 1 µg (0.1 µg/µL). **Digestion**. For the solely trypsin samples, digestion with trypsin (1:100 enzyme:protein ratio; Promega) was carried out overnight at 37 °C. **Recovery of peptides from beads**. For the Sera-Mag™ SP3 bead and ReSYN bead methods, tubes were placed on the magnetic separator and the peptide mixture was carefully pipetted off and dispensed into a fresh microcentrifuge tube. For the glass bead method, tubes were centrifuged (16,000 *g*, 3 min (2x, with tubes spun in between), 4 °C) and the peptide mixture was carefully pipetted off and dispensed into a fresh microcentrifuge tube.

#### Experiment 2

Cell lysates were reduced (5mMDTT, 30 min, 25 °C) and alkylated (5mM iodoacetamide, 30 min, 25 °C in the dark). Proteins were recovered by one of two methods. For the acetone precipitation method, an analogous procedure to Experiment 1 was used up to point of redissolving the pellet. Each pellet was resuspended in the following volumes and buffers: 250 µL of 50mM HEPES for trypsin-only and 125 µL of 50mM HEPES with 1M guanidinium hydrochloride for Lys-C/trypsin. Samples were sonicated and vortexed periodically to redissolve the pellet. For the glass bead method, an analogous procedure to Experiment 1 was used up to point of redistributing the beads. Beads were re-distributed via sonication with the following volumes and buffers: 250 µL of 50mM HEPES for trypsin-only and 125 µL of 50mM HEPES with 1M guanidinium hydrochloride for Lys-C/trypsin. Tubes were centrifuged (16,000 *g*, 3 min (2x, with tubes spun in between), 4 °C) and the peptide mixture was carefully pipetted off and dispensed into a fresh microcentrifuge tube. **Digestion**. For the solely trypsin samples, digestion with trypsin (1:100 enzyme:protein ratio; Promega) was carried out overnight at 37 °C. For the Lys-C/trypsin samples, digestion with Lys-C (1:100 enzyme:protein ratio; Wako) was carried out for 4 h at 37 °C, followed by 1:2 dilution with 50mM HEPES and a secondary digestion with trypsin (1:100 enzyme:protein ratio; Promega) performed overnight at 37 °C.

#### Data acquisition

Assuming 100% recovery, 1 µg of each peptide mixture was added to 200 µL of 0.1% formic acid on a prepared Evotip and run on an Evosep1 LC connected to the Orbitrap*TM* Fusion Lumos MS instrument using 44 min LC-MS gradient in DDA mode as described: the transfer capillary set to 300 °C and 2.2 kV applied to the nanospray needle (Evosep). MS*1* data was acquired in the Orbitrap*TM* with a resolution of 60k, with a max injection time of 20 ms and an AGC target of 1×10^6^, in positive ion mode, with profile spectra, over the mass range 375–1200 m/z. A charge state inclusion of precursors with 2–6^+^ charges was applied with the MIPS mode (Peptide) active, a dynamic exclusion of 15 s, intensity threshold of 5×10^4^, and isolation carried out in the quadrupole with a width of 1.4 Da. For fragmentation, HCD energy of 32% was applied and MS*2* were acquired in the Orbitrap™ with 15k resolution, max injection time of 22 ms and an AGC target of 1×10^6^ in centroid mode.

#### Data analysis

For sample-specific spectral library generation, data was acquired from samples from each condition in data-dependent acquisition (DDA) mode. The data were searched against the human Uniprot database using the Pulsar search engine (Biognosys AG). The following modifications were included in the search: Carbamidomethyl (C) (Fixed) and Oxidation (M)/Acetyl (Protein N-term) (Variable). A maximum of two missed cleavages for trypsin were allowed. The identifications were filtered to satisfy FDR of 1% on peptide and protein level. Protein Group, Peptide and Precursor numbers were reported based on the library generated by the search.

### Lab B

Same as in main methods with differences noted in Fig. 4D.

### Lab C

#### Comparison of SP3 and SP4 sample processing methods

E14 murine embryonic stem cells were lysed in RIPA buffer by pipetting and sonication. The lysates were clarified by centrifugation (20,000 *g*, 10 min, 4 °C) and protein concentrations were determined by BCA assay. Aliquots (n = 5 per experimental condition) corresponding to 50 µg of total protein were removed and diluted (1:1) with 20mM HEPES, pH 8.5 buffer. Reduction with 5mM TCEP final concentration was carried out at 37 °C for 45 min and alkylation with 20mM 2-chloroacetamide (25 °C, 30 min). SP3 and SP4 protocols were carried out as described in the materials and methods section. Following the respective processing methods, rapid digestion buffer (150 µL per sample, Promega VA 1061) was added followed by 5 µg Lys-C/trypsin mixture (Promega VA1061). Protein digestion was carried out at 70 °C with shaking (800 rpm) for 2 h. Samples were removed from the incubator and cooled on ice. Acidification was achieved by addition of 10% TFA (final concentration: 0.25 %) and glass or magnetic beads were removed by centrifugation (20,000 *g*, 5 min, 25 °C). Supernatants were transferred to sample vials and analysed by LC-MS/MS.

#### Liquid chromatography-tandem mass spectrometry (LC-MS/MS) analysis

Sample aliquots corresponding to 1 µg total digest were injected on a U3000 RSLC nano-LC system onto a trapping column (Thermo Acclaim Pepmap 100, 0.1 × 20 mm, 164564) at a flow rate of 8 µL/min with loading buffer (2% acetonitrile, 0.1% TFA). Following valve switch peptides were eluted onto an analytical column (Thermo EASY-Spray™ column, 0.075 × 500 mm, ES803A) by applying a linear multi-step gradient (buffer A: 5% DMSO, 0.1% formic acid; buffer B: 75% acetonitrile, 5% DMSO, 0.1% formic acid) at a flow rate of 250 nL/min and a column temperature of 40 °C: 1% B [0–5 min], 22% B [75 min], 42% B [95 min], 87% B [95.1 min]. The elution gradient was followed by column wash and equilibration steps. The Q Exactive HF-X mass spectrometer was operated in positive ionisation mode at a spray voltage of 1.6 kV. Data-dependent acquisition was carried out with a Top30 method, automatic gain control targets of 3×10^6^ (MS^1^) and 5×10^4^ (MS^2^) ions and maximum accumulation times of 25 ms (MS^1^) and 50 ms (MS^2^), respectively. Dynamic exclusion of fragmented precursors was enabled for 50 s.

#### Data processing and analysis

Raw data files were processed with MaxQuant version 1.6.10.43 and database searches carried out against a Swissprot *Mus musculus* database (version 2020.11.11, 17,056 entries). Settings included trypsin digestion with up to two missed cleavages, and a false discovery rate (FDR) of 1% for peptide spectrum matches and protein identifications. Protein N-terminal acetylation, methionine oxidation and peptide N-terminal glutamine to pyroglutamate conversion were enabled as variable modifications and cysteine carbamidomethylation as a fixed modification. The ‘match between runs’ option was enabled within experimental conditions (SP3 or SP4 digests), with match and alignment time windows of 0.7 and 20 min, respectively.

**Fig. S1.**
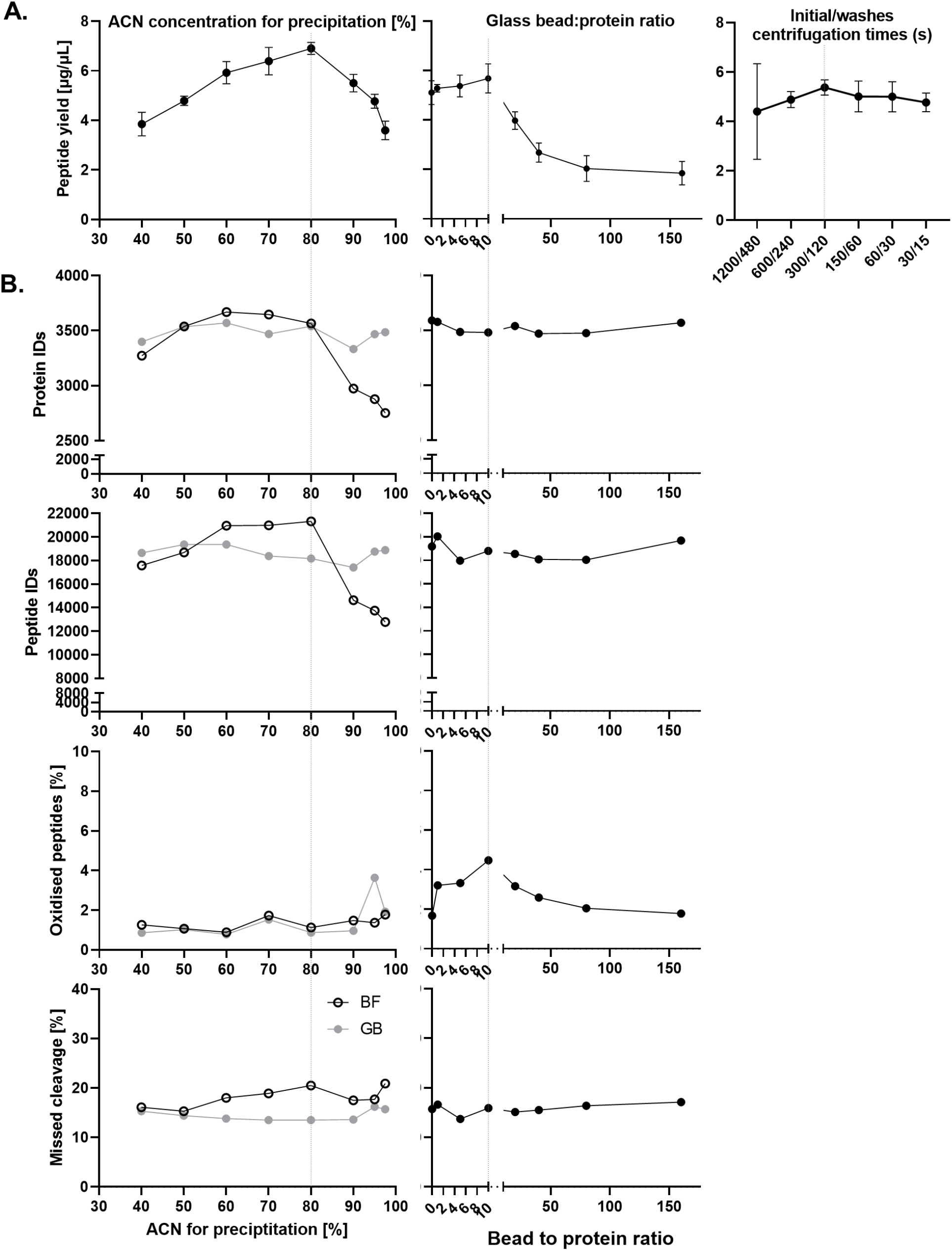
Evaluation of a range of SP4-related variables by peptide quantitation assay (n = 4) and proteomics analysis. **A**. 10 ug of protein was processed by SP4 varying: the initial and post-wash precipitate capture centrifugation times, the glass bead to protein ratio and the total final percentage of ACN in the precipitation step. The digests were measured by peptide quantitation assay. **B**. For proteomics analyses, 10 ug SP4 sample preparations were evaluated varying bead input and ACN concentration with 100 ng equivalent of peptides analysed by LC-MS. Other variables were kept at either 300/120s capture/wash centrifugation steps, 10:1 glass bead to protein ratio and 80 % ACN.

**Fig. S2.**
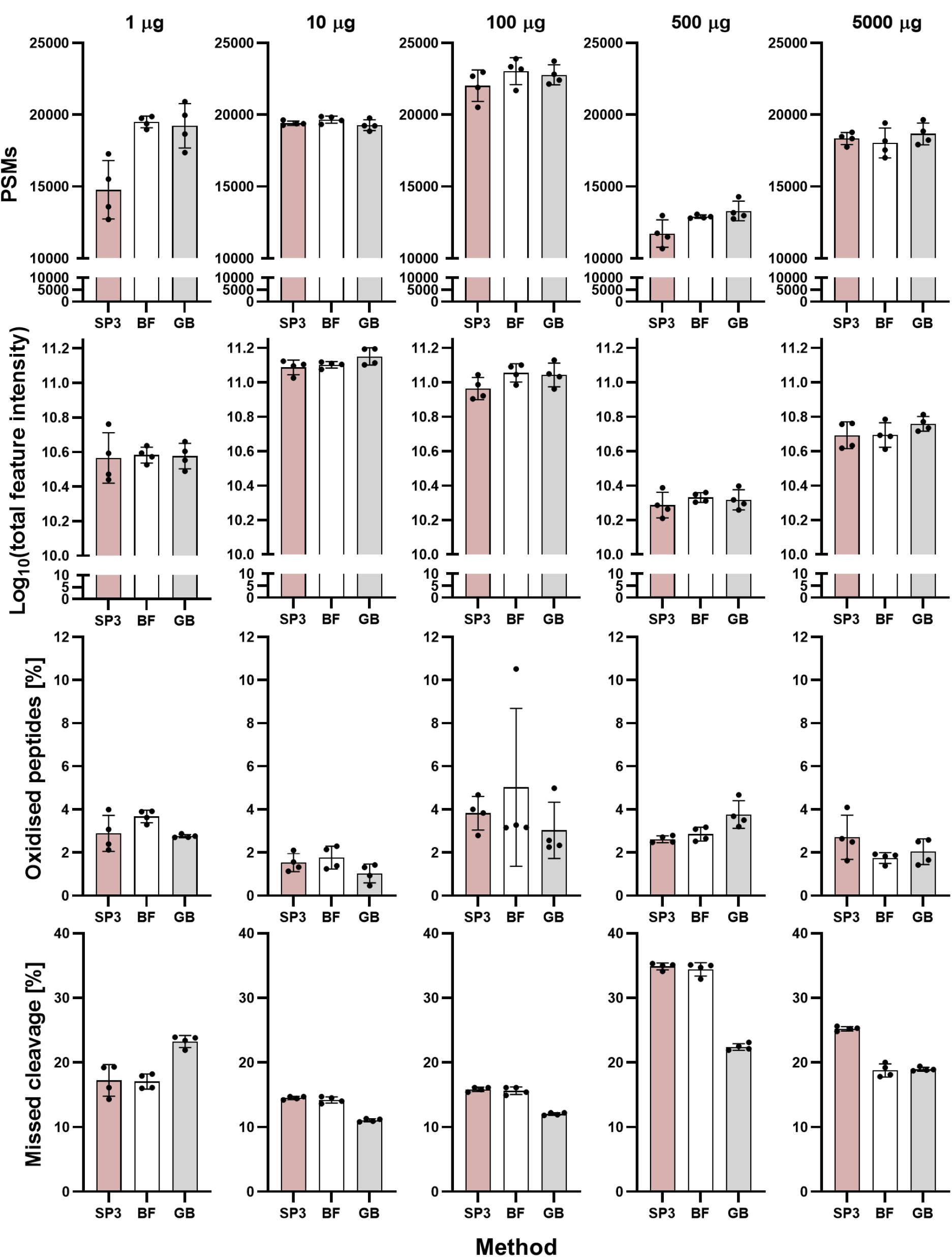
Additional measure of proteome quality and protein recovery for the comparison of SP3 to SP4 with (GB) and without glass beads (bead-free, BF) across a range of protein inputs (see also Fig. 2). PSM = peptide spectrum match.

**Fig. S3.**
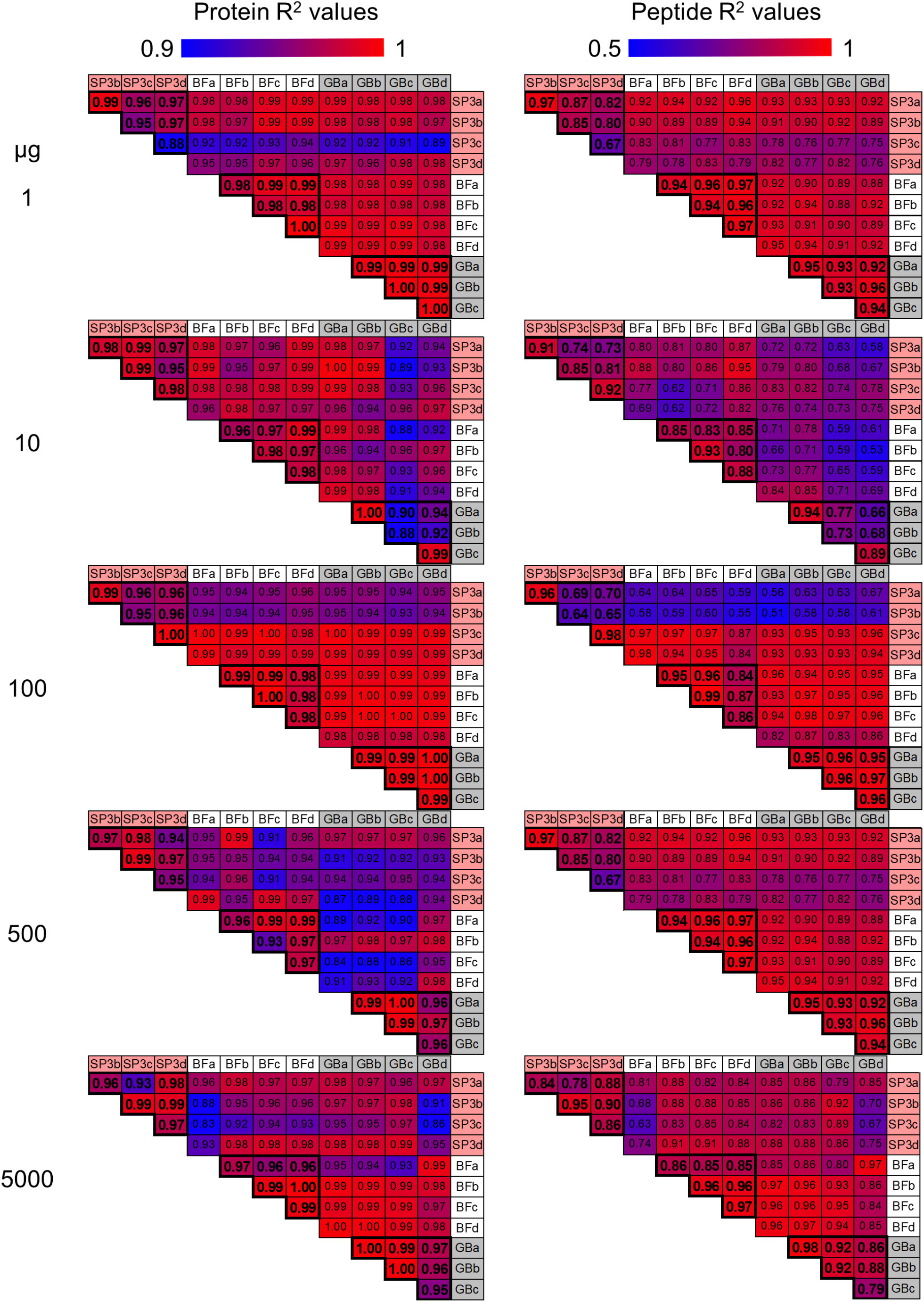
Protein and peptide LFQ R^2^ values of recovery for SP3 to SP4, with (GB) and without glass beads (bead-free, BF), across a range of protein inputs (see also Fig. 2).

**Fig. S4.**
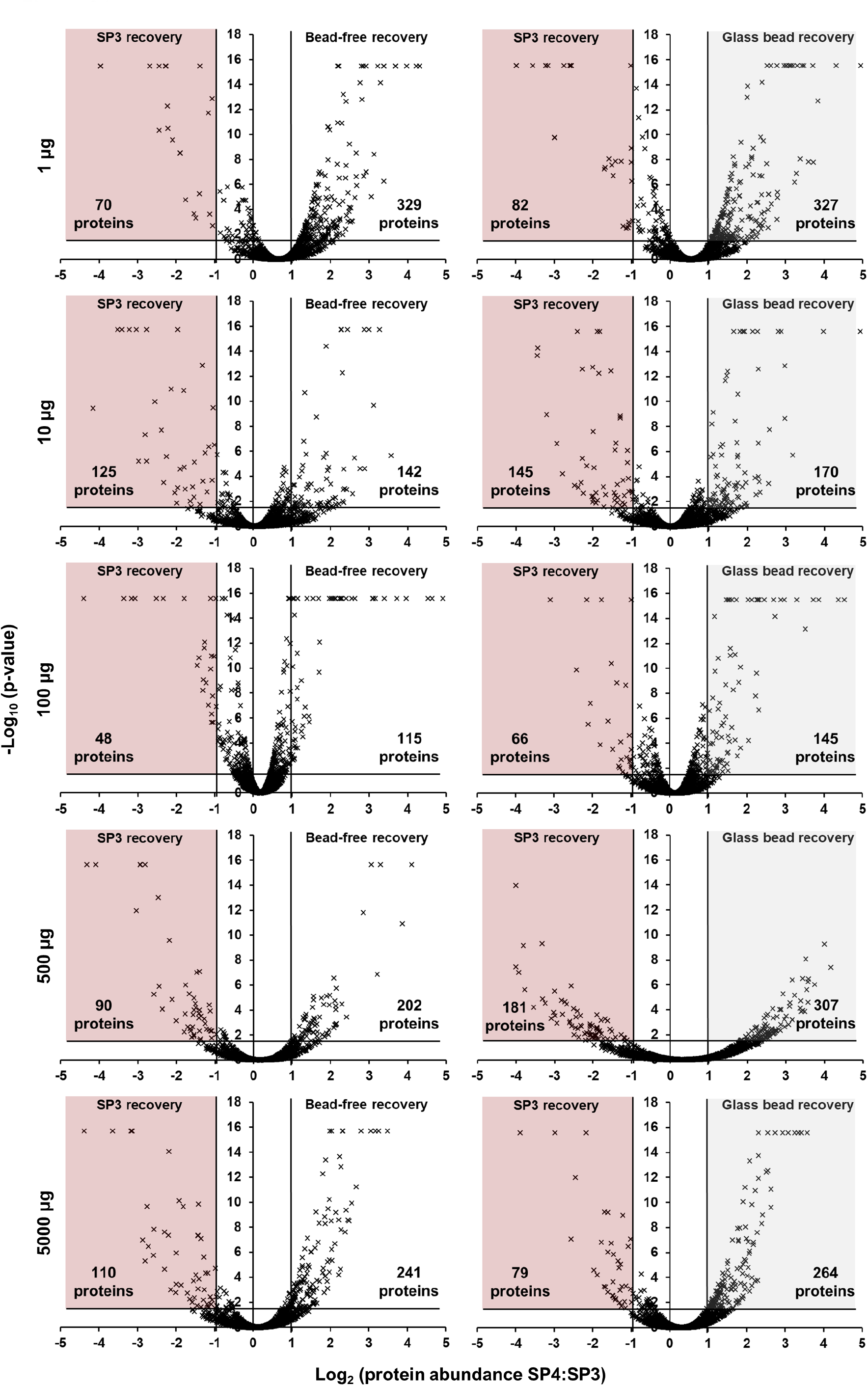
Protein recovery observed to be significantly more effective by SP4 variants versus SP3, summarised in Fig. 2.

**Fig. S5.**
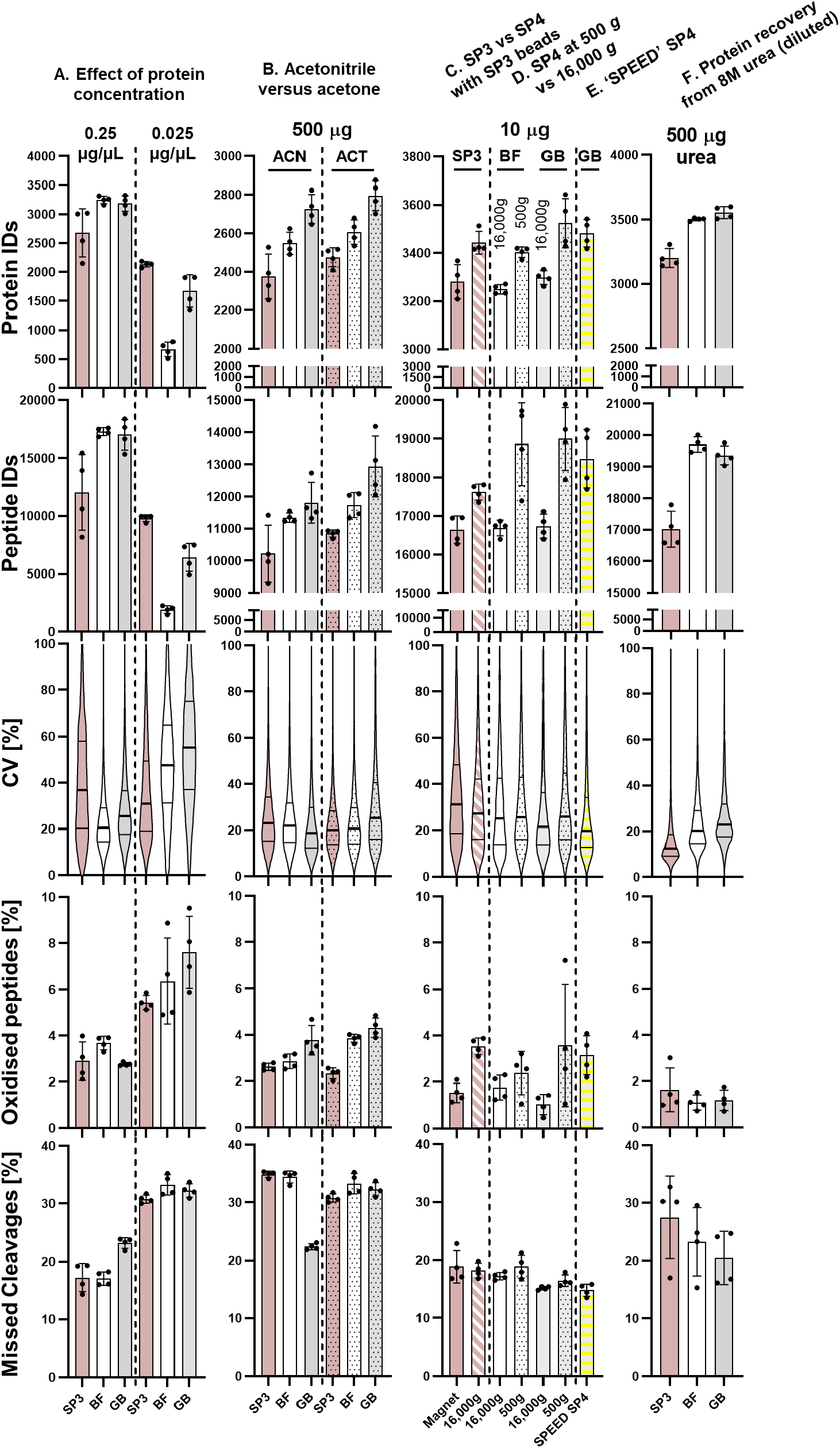
Additional experiments exploring the mechanism and potential of bead-free (BF) and glass bead (GB) SP4. Protein numbers, peptide numbers, peptide CVs, oxidised peptides, and missed cleavage rates among peptide spectrum matches are detailed for the 4 procedural replicates. **A**. 1 µg SP3 and SP4 preparations were conducted using two initial protein concentrations: 250 ng/µL (1 µg in 4 µL volume, including beads) and 25 ng/µL (1 µg in 20 µL volume, including beads). **B**. 500 µg SP3 and SP4 preparations were conducted using acetonitrile (ACN) and acetone (ACT) as the denaturing solvent. **C**. SP3 was compared with SP4 using SP3 carboxylate magnetic beads to confirm that centrifugation recovered more protein than the use of a magnet. **D**. BF and GB SP4 variants were tested at 500 *g* (*vs*. 16,000 *g* adopted in all other experiments) for the potential to expand their compatibility with larger volume and plate-based preparations. **E**. Cells were lysed by ‘SPEED’ (<u>S</u>ample <u>P</u>reparation by <u>E</u>asy <u>E</u>xtraction and <u>D</u>igestion) method using 100% TFA and neutralised with Tris base before being subjected to SP4 with the inclusion of glass beads. **F**. 500 µg of protein was processed by SP3 to SP4 using 8M urea as the lysis buffer, across a range of measures of protein recovery and proteome quality. Lysate was diluted to 2M urea prior to addition of beads and ACN.

**Fig. S6.**
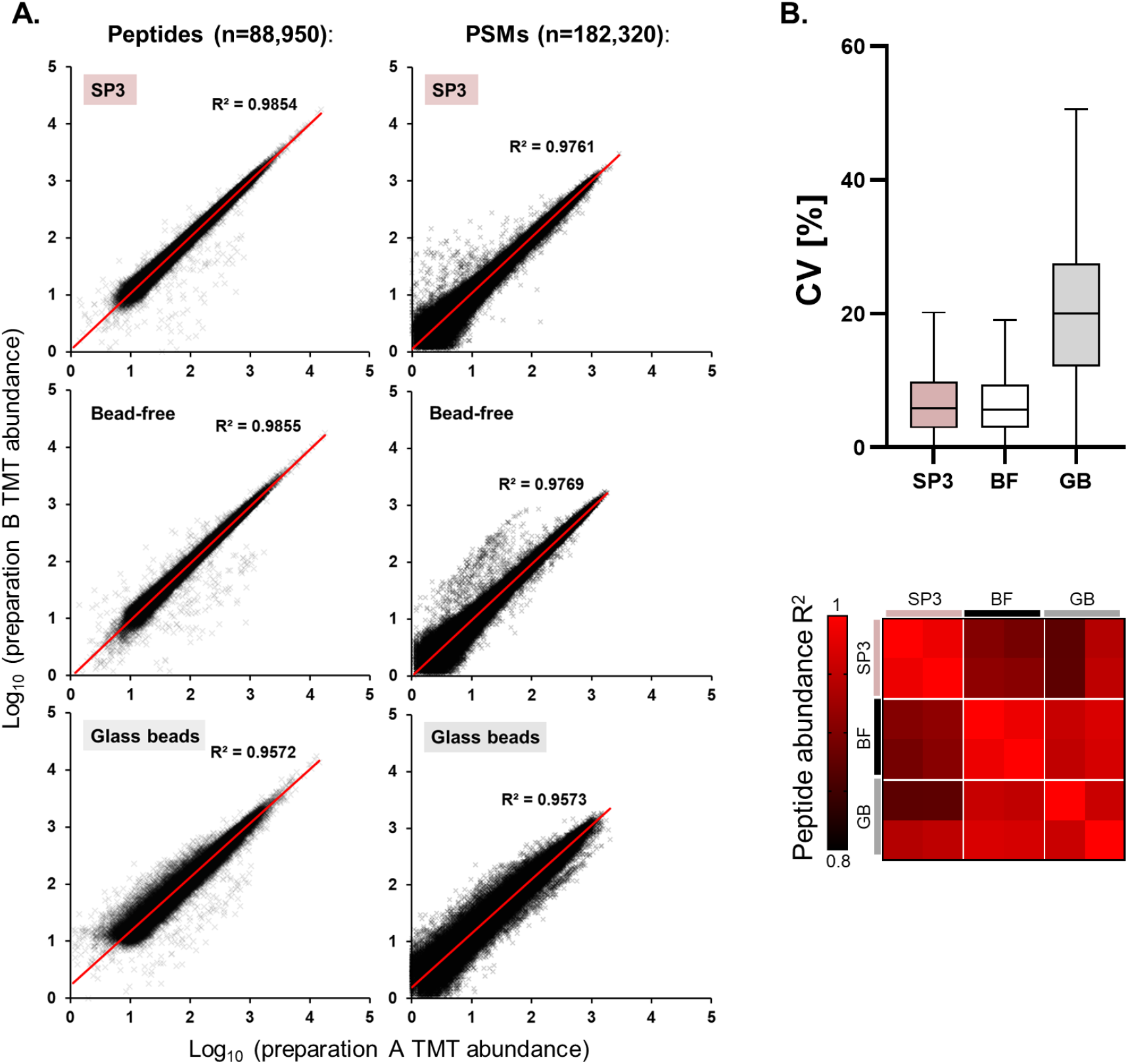
Additional measures of quantitative proteome quality for comparison of SP3 to SP4 with and without glass beads using TMT 6-plex, as summarised in Fig. 3. **A**. Correlation between TMT-measure peptide and PSM abundances for sample preparation method replicates. **B**. Coefficients of varation (CV) and R^2^ values for peptide quantification between the method replicates.

**Fig. S7.**
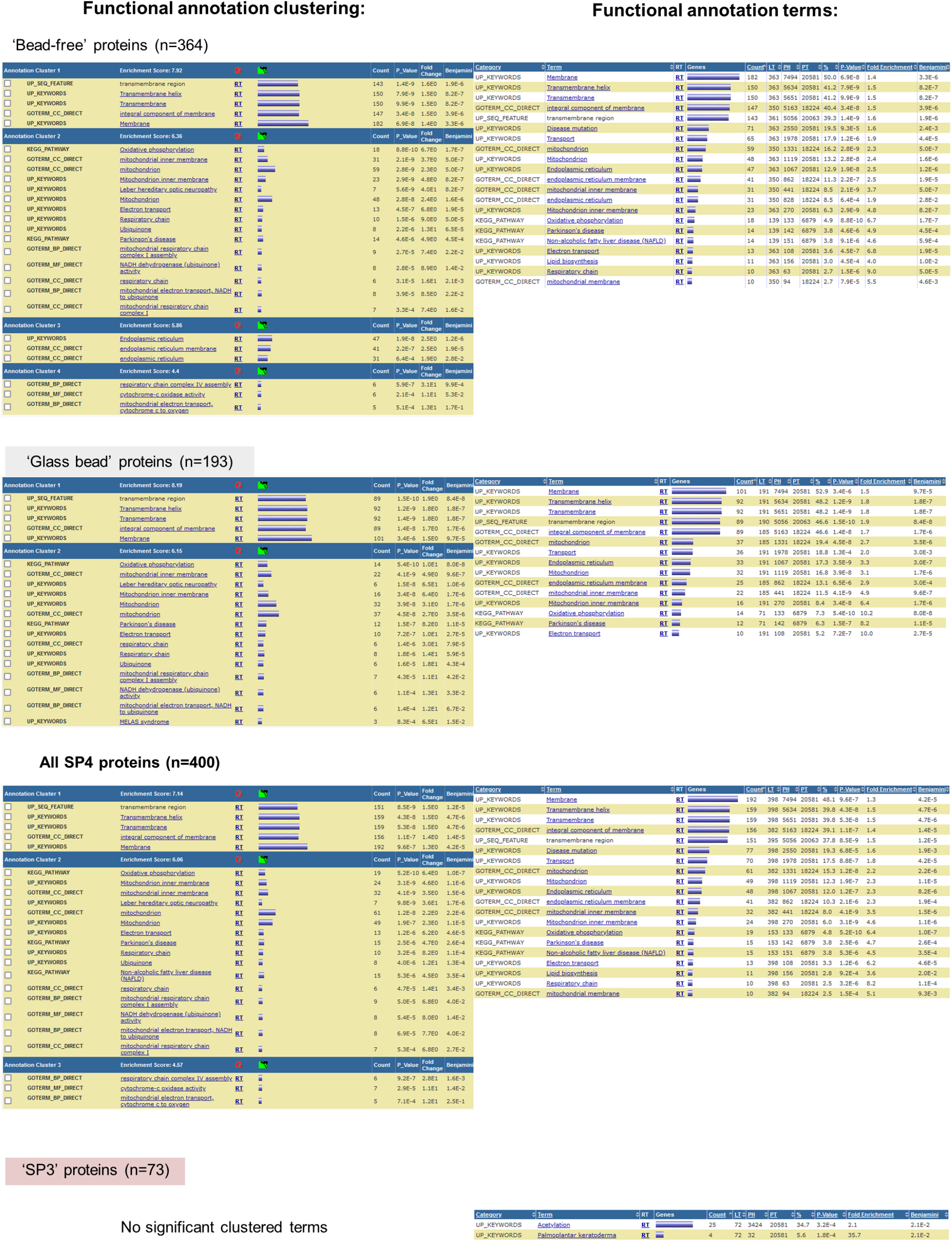
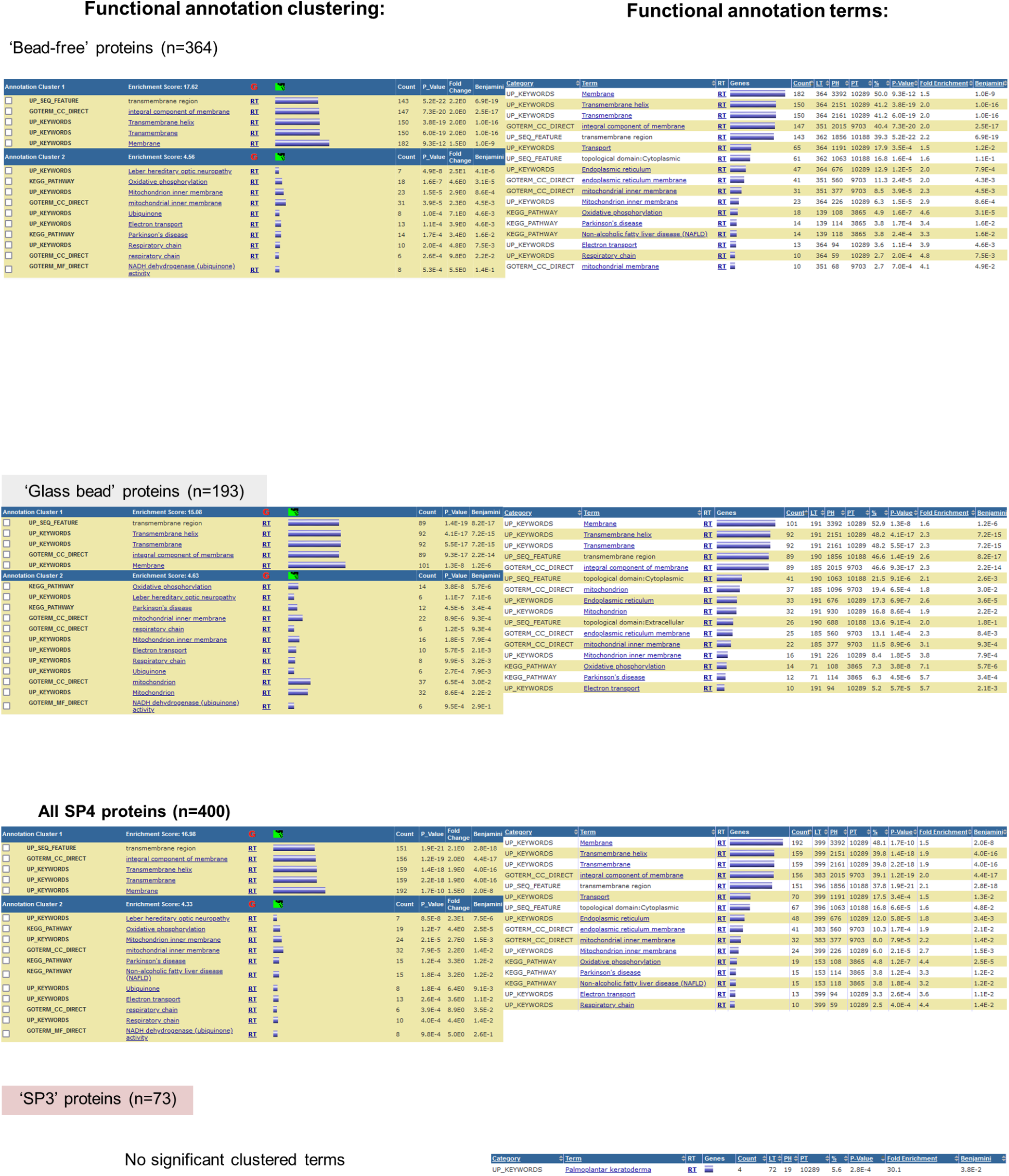
DAVID-derived term enrichment and clustering for those proteins observed more significantly recovered by SP3 and SP4 by TMT quantification. Terms were filtered to include those with a significant enrichment before (p < 0.001) and after (p < 0.05) Benjamini correction, and to include at least 10 terms (SP4 only, for clarity). Both the TMT proteome-identified proteins and the whole human Swissprot proteome were used as background, to highlight the greater significance of over-representation amongst proteomics-identified proteins.

## SP4 (Solvent precipitation SP3) protocol

### Glass bead preparation (optional)

- 9-13 μm glass spheres/beads (e.g., https://www.sigmaaldrich.com/catalog/product/aldrich/440345) Glass beads broadly improved recovery, digestion efficiency and reproducibility, but are not essential
- Suspend 100 mg in 1 mL of ultrapure water, vortex until suspended fully, and pellet at >500 *g*, 1 min. Of note: approximately 50 % of the beads are buoyant, and will not pellet, and should be removed over the course of these wash steps. Additionally, small amounts of metal in the beads can be removed by magnet or acid wash but had no effect on the performance of the beads.
- Resuspend, vortex and wash with >1 mL of: 100% acetonitrile (ACN) (1x), 100 mM ABC* (1x), and ultrapure water (2x). * or equivalent digestion buffer.

Then either:

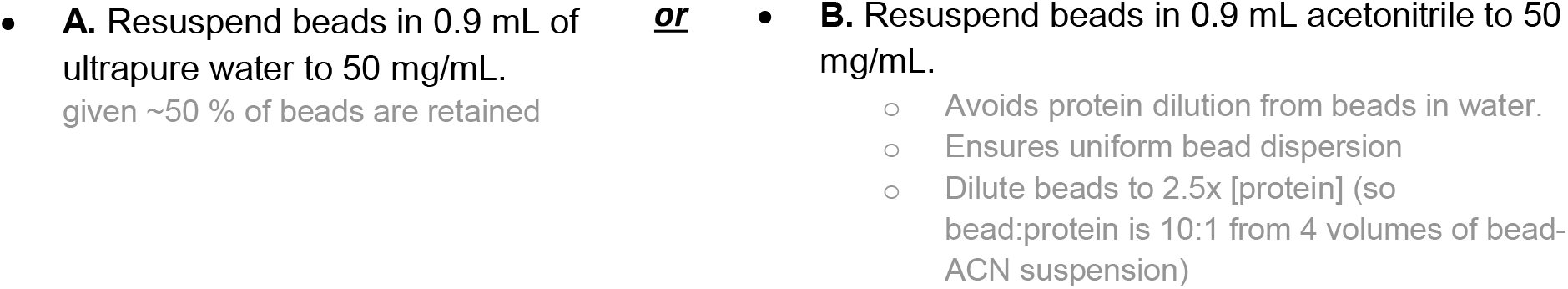

This will be sufficient to prepare 50 mg of protein—excess can be stored at 4°C.

**(with 0**.**2% sodium azide, if in water)**

### Lysate/protein solution prep recommendations

- SP4 is broadly compatible with the majority of lysis buffers as for SP3 or acetone precipitation Tested with:
  - 5% total detergent ‘SP3 lysis buffer’ (50 mM HEPES pH 8, 1% SDS, 1% Triton X-100, 1% NP-40, 1% Tween 20, 1% Na deoxycholate, 50 mM NaCl, 5 mM EDTA and 1% (w/v) glycerol, 1 × cOmplete™ protease inhibitor)
  - 8M urea (diluted to 2 M prior to ACN addition)
  - TFA/TRIS diluted 1:1 with water as described for the ‘SPEED’ method
- For best results with SP4, protein concentration should be as high as possible (0.25–5 µg/µL).
  - For lower concentrations or where highest possible recovery is required, longer precipitation reactions, pre-chilled CAN, and centrifugation at 4°C may help yields.
- DNA shearing (e.g., by sonication), protease inhibitors, & lysate clearance are recommended.

### SP4 protocol recommendations

- The use of the smallest possible tube will help create a denser pellet, e.g., 500 µL tube for samples of less than 50 µL.
- Liquids should be kept low in the tube, with losses/contamination possible from tube walls/lid.
- Set vortex to <500rpm for very gentle mixing.
- Pipette ACN directly into the sample to ensure rapid mixing, but do not touch the ACN/sample mix with the tip.
- Use the tube hinge to orientate the location of the pellet (fixed angle rotors).
  - Initially orientate the tube hinge inwards during the pellet precipitation and turn 180° after 2.5 min will give a denser pellet and less risk of loss from fragile wall adhesion.
- During wash removals, avoid touching the tube walls with the tip as precipitation may occur on them, pipette slowly and avoid agitating the pellet.
- If adding beads, ensure uniform suspension by pipetting up and down at least once between additions.

### SP4 Protocol

1. Aliquot reduced/alkylated protein mixture/lysate into a fresh lobind-type microcentrifuge tube.
  - Volumes and conditions are given for the example of 10 µg protein in 10 µL of 1 µg/µL lysate.
2. Options (choose one):

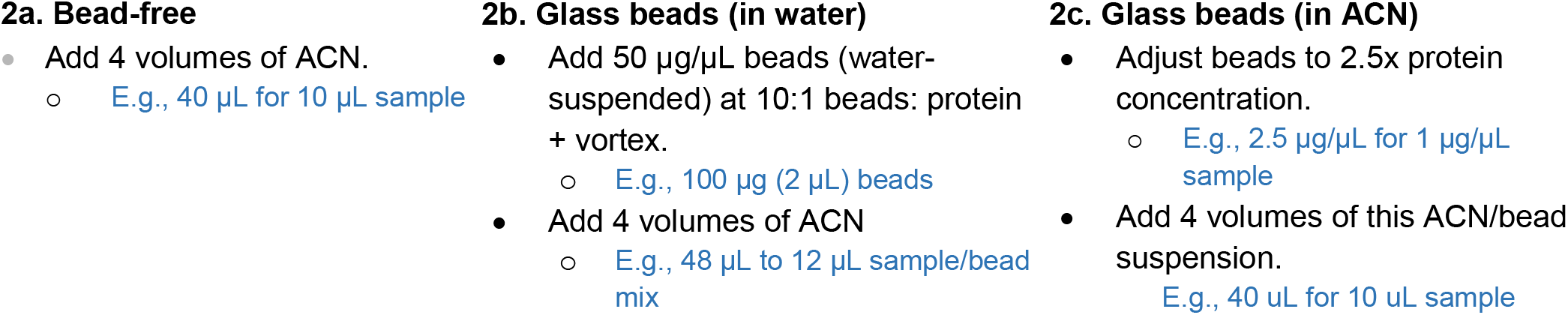
3. Gently vortex on a low setting to mix for 5 sec (do not pipette mix).
4. Centrifuge for 5 min at 500–16,000 *g*.
5. Remove supernatant by pipetting slowly and remove 90–95%. Avoid disturbing beads/pellet. E.g., leaving <10 µL
6. Wash with 80% ethanol, volume >= 1.5x total precipitation volume.
  - Pipette gently down the side opposite the hinge/pellet to avoid disturbance, do not vortex/resuspend. E.g., >150 µL for washes
7. Centrifuge for 2 min at 16,000 *g*.
8. Remove 90-95% of wash. E.g., leaving <10 µL
9. Repeat wash steps for a total of 3 washes.
10. Remove >=95% of final wash.
  - For larger volumes a final 2 min spin will help with removal of excess wash. E.g., leaving <5 µL
11. Add preferred digestion buffer, e.g., 20-100 mM ABC or TEAB.
12. Add preferred digestion enzyme, e.g., trypsin/Lys-C at a 1:25 to 1:100 enzyme:protein ratio.
  - A digestion buffer/enzyme master mix will reduce variability and simplify pipetting - keep on ice.
  - Use a volume equivalent to ∼0.5-2x the total precipitation volume. E.g., 25-100 µL for 50 µL precipitation reaction
  - In-bath sonication (5-10 min) can help to disrupt the pellet and increase surface area.
  - Pipette mixing may cause losses especially once trypsin is added.
  - Larger bead-free pellets may require additional agitation to resuspend but keep sample low in tube. 18 h digestion consistently worked without pellet resuspension.
13. Incubate in a thermomixer at 1000 rpm at desired conditions, e.g., for 18 h at 37 °C.
  - Use of beads enhanced digestion and was compatible with a 2 h, 70°C incubation using a rapid digestion buffer.

### Peptide collection

- Centrifuge the peptide mixture at 500–16000 *g* for 2 min & collect peptide supernatant.
- For maximum recovery, rinse pellet in an equal volume of digestion buffer added above.
  - A final centrifugation step may be required to ensure no beads are carried over.
- Peptides solution at this stage is clean enough for:
  - Acidified for direct LC-MS injection.
  - Dried by vacuum concentration to provide near-pure peptides.

